# Dual cellular targeting of complement receptor 2 (CR2/CD21) synergistically suppresses autoreactive germinal center activity and autoimmunity in lupus

**DOI:** 10.64898/2026.11.12.761594

**Authors:** Ajitanuj Rattan, Danni Yi-Dan Zhu, Kristine Oleinika, Minghe Ma, Michael C. Carroll

## Abstract

Pathogenic germinal center (GC) activity persists in systemic lupus erythematosus (SLE) because current B cell-targeting therapies do not consistently eliminate the resilient GC niche that sustains autoimmunity. Complement receptor 2 (CR2/CD21) presents a unique bicellular therapeutic target to impede GC-associated autoimmune disease. For B cells, it has been previously shown to lower the threshold for B cell activation, and for follicular dendritic cells (FDCs), it sustains the GC response. Using a CD21-blocking antibody in a spontaneous lupus-prone mouse model, we demonstrate that dual CD21 blockade on both B cells and FDCs synergistically inhibited GC B cell responses and suppressed autoantibody production, memory recall, and renal antibody deposition. Mechanistically, B cell-specific blockade impaired GC entry by limiting initial activation, whereas FDC-specific blockade disrupted GC architecture and curtailed ongoing responses. Further, dual blockade uniquely reshaped immunological memory by promoting accumulation of a CD21^low/-^CD38^+^ memory B cell subset with attenuated antibody-secreting cell (ASC) recall potential. These findings identify CD21 blockade as a precision immunomodulatory strategy capable of suppressing pathogenic humoral immunity without complete B cell depletion.

**One sentence summary:** Dual CD21 blockade on B cells and follicular dendritic cells synergistically suppresses autoreactive germinal center responses and reshapes B cell memory in a lupus mouse model.

## Introduction

Systemic lupus erythematosus (SLE) is a chronic autoimmune disease characterized by the production of pathogenic autoantibodies, immune complex (IC) deposition, and progressive multiorgan inflammation (1–3). A critical feature of SLE pathogenesis is the formation of chronic GCs within secondary lymphoid tissues, where activated B cells undergo clonal expansion, somatic hypermutation, affinity-based selection, and differentiation into memory B cells (MBCs) and ASCs (4–8). Within these chronic autoreactive GCs, self-reactive B cells undergo iterative diversification and selection, leading to maturation of existing autoreactivity and progressive epitope spreading toward additional self-antigens (9). Moreover, spontaneous autoreactive GCs are dynamic structures that can rapidly recruit otherwise naïve wild-type B cells into the autoreactive GC response, enabling newly recruited clones to expand, persist, and contribute to diversification of the autoimmune repertoire (10). Thus, autoreactive GC responses represent an evolving reservoir capable of continually expanding both the magnitude and antigenic breadth of humoral autoimmunity (9–13).

Current B cell-directed therapies represent an important advancement in SLE management, but the removal of circulating B cells does not necessarily dismantle the GCs maintaining or regenerating autoreactive humoral responses (23,24). Belimumab, which neutralizes B cell-activating factor (BAFF), can reduce the survival and replenishment of circulating B cells, whereas anifrolumab-fnia, an antagonist of the type I interferon receptor, suppresses inflammatory signaling that promotes autoreactive B cell activation (15–17). However, neither therapy directly eliminates established autoreactive GCs nor disrupts the specialized cellular niches that support them. The GC microenvironment is maintained by specialized interactions among GC B cells, FDCs, and T follicular helper (TfH) cells, wherein FDCs sustain antigen availability, TfH cells promote affinity-based selection, and reciprocal signaling supports GC B cell survival (14,18,19). Consequently, GC-resident B cells may persist despite systemic reductions in B-cell survival signals or inflammatory activity, allowing autoreactive humoral responses to continue or re-emerge. Consistently, anti-CD20-mediated B cell depletion has not uniformly achieved clinical benefit in randomized SLE trials, including trials in both non-renal and renal disease (20,21), and lymphoid-tissue analyses following rituximab therapy have demonstrated active GC reactions despite profound depletion of circulating B cells (22).

Chimeric antigen receptor (CAR) T cell therapy targeting CD19, which reaches a broader population of B cells than CD20, has produced drug-free remission in severe refractory SLE cohorts, including sustained remission after B cell reconstitution, suggesting that sufficiently deep disruption of the autoreactive B cell compartment can functionally reset humoral autoimmunity (25,26). Nevertheless, CAR T cell therapy requires lymphodepleting conditioning and specialized cellular-therapy infrastructure and can be associated with cytokine release syndrome, infectious complications, and a period of B cell aplasia. Therefore, its long-term durability and optimal positioning within SLE therapy continue to be defined (25,26). There is a need for complementary therapeutic strategies capable of selectively disrupting pathogenic autoreactive programs and their supporting microenvironments without requiring broad lineage ablation.

CD21 occupies a particularly attractive position within this therapeutic landscape because it regulates humoral immunity at both the B cell and FDC levels. On mature B cells, CD21 associates with CD19 and CD81 to form the B cell co-receptor complex, enabling complement-tagged antigen to amplify B cell receptor (BCR) signaling and markedly lower the threshold required for B cell activation (27,28). Complement regulation of B cell activation through CR2/CD21 is especially important when antigen availability or BCR affinity is limiting, conditions that may be highly relevant during early recruitment of autoreactive B cells (27–31). On FDCs, expression of CR1/CR2 is required for efficient retention of immune complexes and sustained antigen-specific humoral responses (35). FDCs acquire complement-opsonized immune complexes through complement receptors, internalize them into nondegradative compartments, and repeatedly recycle intact antigen to the cell surface, thereby maintaining an antigen reservoir accessible to GC B cells during selection and affinity maturation (18,36). In autoimmunity, FDCs can capture complement-opsonized ribonucleoprotein (RNP) ICs through CD21, leading to endosomal TLR7 activation, IRF5-dependent type I interferon production, and promotion of autoreactive GC responses (37). Thus, FDC-associated CD21 is not simply an antigen-retention receptor but a component of an active signaling niche capable of sustaining chronic autoreactive B cell responses.

Importantly, CD21 is pharmacologically targetable. Structure-guided studies identified inhibitory anti-human CD21 monoclonal antibodies capable of directly blocking the CR2/CD21-C3d interaction on B cells (32). Subsequent *in vivo* studies using human CR2/CD21 transgenic mice demonstrated that antagonistic CD21 antibodies suppress complement-dependent B cell activation and antibody responses without requiring B cell depletion (33). Receptor-targeting antibodies have also been developed for chronic manipulation of murine CR1/CR2 function (34). Recent work by Zhu *et al.* demonstrated that continuous CD21-complement engagement precedes TLR7-dependent extrafollicular differentiation of naïve autoreactive B cells in the 564Igi mouse model and showed that functional blockade of human CD21 suppresses progression toward the autoreactive ASC program (39). However, the GC-dependent and extrafollicular pathways function as complementary sources of autoreactive effector cells in SLE, and the contribution of CD21 to established lupus-associated GC responses and their supporting microenvironment has remained unresolved (40).

We therefore hypothesized that therapeutic CD21 blockade could simultaneously target two mechanistically distinct cellular compartments of autoreactive humoral immunity: 1) activation-dependent recruitment of autoreactive B cells through the B cell co-receptor and 2) maintenance of the pathogenic GC niche through FDC-associated complement receptors. To test this hypothesis, we used a CD21-blocking monoclonal antibody together with human CD21-transgenic mice (hCD21^+^mCD21^-/-^; hCD21*tg*) (33) to generate compartment-specific mixed bone marrow (BM) chimeras. We also made adoptive transfer models using the 564Igi mouse model, in which B cells carry a genetically engineered BCR targeting RNP-associated self-antigens (38). In these mixed BM chimeras, 564Igi-derived B cells are reconstituted at a defined ratio alongside wild-type or hCD21tg donor B cells and must therefore compete with non-autoreactive B cells for GC entry; despite this competitive disadvantage, 564Igi-derived B cells reliably generate a quantifiable, reproducible autoreactive GC and class-switched autoantibody response within the post-reconstitution timeframe used throughout this study (6–8 weeks), recapitulating key features of autoreactive humoral immunity, including spontaneous GC formation, recruitment of naïve B cells into ongoing autoreactive responses, class-switched autoantibody production, and renal immune-complex deposition (9,10,38,39). We identify CD21 blockade as a precision immunomodulatory strategy capable of suppressing pathogenic humoral immunity without complete B cell depletion.

## Results

### Anti-CD21 antibody achieves efficient in vivo receptor saturation blockade

To achieve compartment-specific CD21 blockade, we first tested the efficacy of a function-blocking anti-human CD21 monoclonal antibody (anti-hCD21) to saturate CD21 receptors on B cells in a dose-dependent manner, by administering single injections of 5, 10, and 20 μg antibody in hCD21*tg* mice (**Fig. S1A**). Mice were bled every day for five days to measure receptor saturation on B cells, frequency of unoccupied hCD21 receptors, and blockade efficiency. A single injection of 20 μg of antibody achieved 85–97% receptor blockade and receptor saturation for 3 days, as detected with the same blocking antibody, clone 658, conjugated to Alexa Fluor-647 (AF647) fluorochrome (**Fig. S1B**).

We then administered 3 injections of 20 μg of blocking Ab to hCD21*tg* mice, and mice received injections every 4 days to keep the receptor occupied for 10 days (**Fig**. **1A**). We measured available (unoccupied) hCD21 by staining B cells with two anti-hCD21 antibodies recognizing distinct epitopes, and quantified blockade efficiency by receptor occupancy (RO) assay. The anti-hCD21 antibody efficiently blocked hCD21 on both B cells and FDCs. Staining with the non-blocking clone Bu32 gave comparable signal in isotype control and anti-hCD21antibody treated animals, indicating that the receptor remained on the cell surface rather than being internalized or shed, and that the loss of blocking-clone signal reflected epitope occupancy (**Fig. 1A, right panel**). Isotype control antibody administered across the same dose range did not reduce blocking-clone staining, confirming that loss of available hCD21 was specific to the CD21-targeting antibody rather than a nonspecific antibody effect (****p < 0.0001, isotype control versus anti-hCD21; **Fig. 1A**). Blocking efficiency was further confirmed by immunofluorescence on B cells (**Fig. 1B**) and FDCs (**Fig. S1C**). FDC staining was performed in B6 WT + 564Igi → irradiated hCD21*tg* chimeras (**Fig. 1C**, *FDC blockade*); because donor B cells lack hCD21 and FDCs are radioresistant and therefore host-derived, follicular hCD21 signal in these mice is attributable specifically to FDCs. Based on this data, we decided to perform 3 or more injections of 20 μg blocking antibody for the rest of the experiments. Total splenic B cell numbers were comparable between anti-hCD21 antibody-, isotype control-, and PBS-treated mice **(Fig. S1D**), confirming that CD21 blockade acts through receptor occupancy rather than depletion of the targeted cell populations.

**Figure 1.**
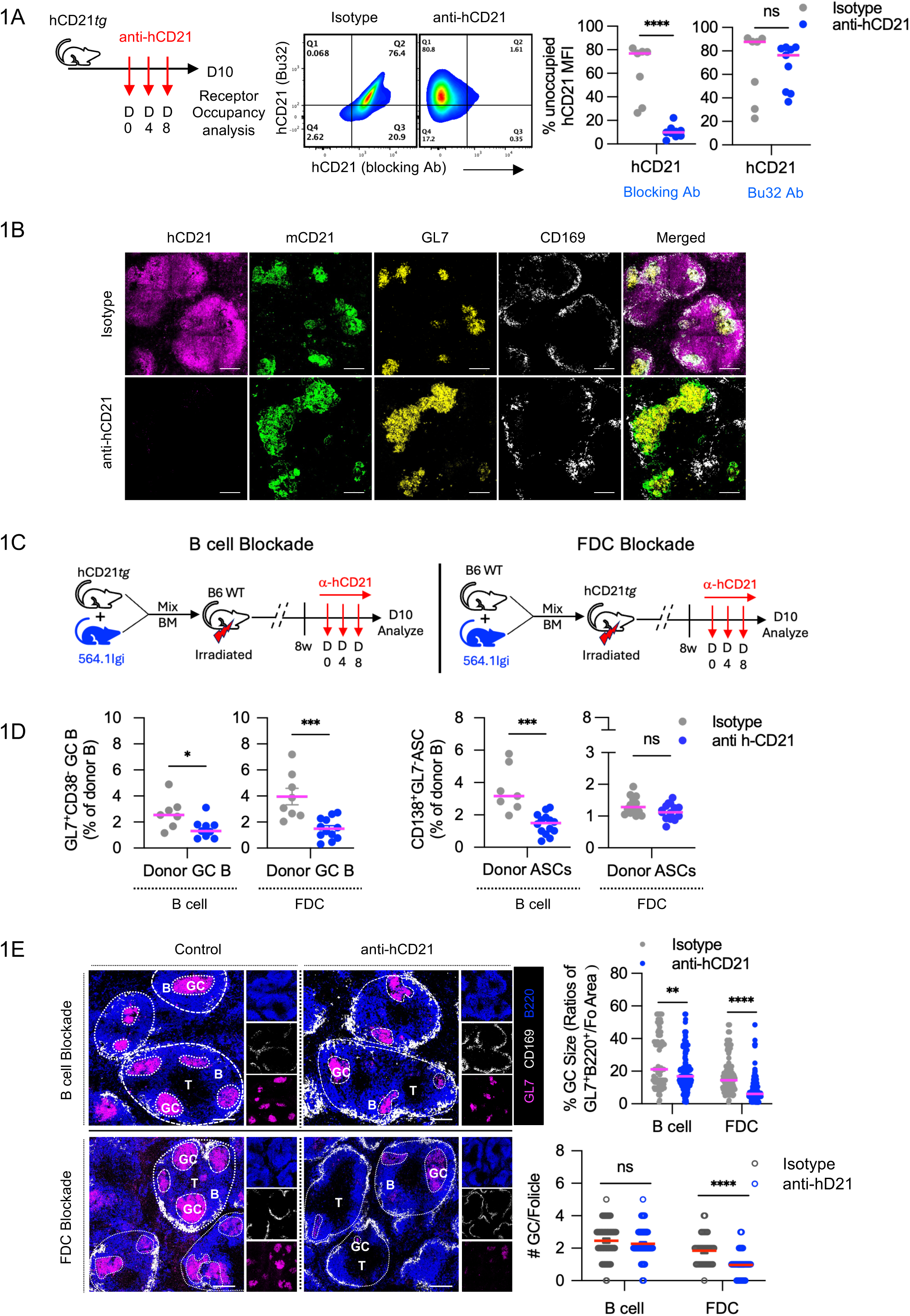
CD21 blockade on B cells and FDCs differentially impacts donor-derived GC B cell and ASC responses in mixed BM chimeras. **A)** Experimental setup to validate efficiency of 20 μg anti-hCD21 blocking antibody administered into hCD21*tg* mice at D0, D4 and D8. Representative flow plots show hCD21 detected with a non-blocking antibody (clone Bu32, y-axis) versus the blocking antibody (x-axis) in isotype and anti-hCD21-treated mice, with quantification of % unoccupied hCD21 for each detection antibody. **B)** Representative immunofluorescence confirming *in vivo* blockade of hCD21 on B cells (magenta) and mCD21 on FDCs (green), with GL7 (yellow) and CD169 (white), in isotype versus anti-hCD21-treated mice. Scale bars = 100 μm. **D) C)** Experimental design for compartment-specific mixed BM chimeras: B cell blockade (hCD21*tg*+564.1Igi BM into B6 WT recipients; left) and FDC blockade (B6 WT+564.1Igi BM into hCD21*tg* recipients; right); both treated with anti-hCD21 antibody (D0, D4, D8) and analyzed at D10. Control mice received equal dose of isotype antibody **D)** Frequency of donor-derived (non-564Igi-origin) GC B cells (GL7^+^CD38^-^) and ASCs (CD138^+^GL7^-^) in B cell-blockade and FDC-blockade chimeras, isotype versus anti-hCD21-treated mice. **E)** Representative immunofluorescence (B220, CD169, GL7) of splenic follicles from B cell-and FDC-blockade chimeras, with quantification of GC size (ratio of GL7^+^B220^+^ area to CD169^+^ follicle area) and GC number per follicle. Data represent mean ± SEM of two independent experiments. *p < 0.05, **p < 0.01, ***p < 0.001, ****p < 0.0001 (Mann–Whitney, Prism v10.1). *ns*, not significant.

### CD21 blockade on B cells and FDCs differentially impacts donor-derived ASC and GC B cell responses

Next, to evaluate the effect of compartment -specific CD21 blockade on total GC B (CD138^-^GL7^+^) cells and ASC (CD138^+^GL7^-^) response, we generated mixed BM chimeras using hCD21tg and 564.1Igi lupus-prone mice (see schematic **Fig. 1C**). To selectively target CD21 on B cells, BM from hCD21tg and 564.1Igi mice was transferred into irradiated B6 WT recipients, generating chimeras with hCD21-expressing donor-derived B cells and WT follicular dendritic cells (FDCs). Conversely, to target CD21 on FDCs, BM from B6 WT and 564.1Igi mice was transferred into irradiated hCD21tg recipients, generating chimeras with WT donor-derived B cells and hCD21-expressing FDCs. In both configurations, 564.1Igi-derived B cells established an autoreactive environment, allowing us to examine the contribution of CD21 in distinct cellular compartments to ongoing autoimmune responses.

Donor derived B cells (CD45.2; hCD21*tg* or B6 WT derived) were distinguished from 564.1Igi B cells using congenic gating for downstream analysis (**Fig. S2A**). Compared to CD21 blockade on hCD21*tg* B cells, where the reduction in GC B cell response was moderate, CD21 blockade on hCD21*tg* FDCs markedly reduced GC B cell response (**Fig. 1D; *left panel,* Fig. S2B).** In contrast, donor-derived ASC frequencies were significantly reduced following B cell-associated CD21 blockade but were not significantly altered by FDC-associated CD21 blockade (**Fig. 1D; *right panel,* Fig. S2C**). This differential effect is consistent with the contribution of CD21-dependent B cell activation to early ASC generation through a GC-independent pathway, as previously reported (39). Consistent with the pronounced reduction in GC B cell responses, histological analysis revealed that FDC-associated CD21 blockade significantly reduced both GC size and the number of GCs per follicle, whereas B cell-associated blockade reduced GC size without significantly altering GC number (**Fig. 1E**). Together, these observations reveal distinct, yet complementary roles for CD21 in autoreactive humoral responses: B cell-associated CD21 preferentially supports ASC responses, whereas FDC-associated CD21 plays a dominant role in maintaining GC B cell responses and GC architecture.

### Dual CD21 blockade synergistically suppresses autoreactive GC responses and autoimmunity

It is clear from these results (**Fig. 1**) that CD21 blockade has a differential effect on donor-derived total ASC and GC B cell responses based on the site of intervention. We therefore investigated whether simultaneous blockade of CD21 on both B cells and FDCs could overcome this differential effect and maximally suppress both GC B cell and ASC responses.

To achieve simultaneous dual CD21 blockade on both B cells and FDCs, we constructed mixed BM chimeras of hCD21*tg* and 564.1Igi lupus mice in irradiated hCD21*tg* recipient mice (see schematic **Fig. 2A**). In these dual-blockade chimeras, we observed synergistic suppression, achieving a greater reduction in both total GC B cell numbers and ASC frequency relative to isotype controls, (**Figs. 2B, S3A-S3C**) than did the single-compartment targeting intervention. We found that while dual CD21 blockade efficiently reduced GC participation and ASC generation by autoreactive donor B cells, no significant difference was observed in donor-derived (IgG2c) anti-ssDNA serum autoantibody levels (**Fig. 2C**), indicating that this short-course regimen (three injections) was sufficient to suppress GC participation and ASC generation at the cellular level but insufficient, within this timeframe, to produce a measurable systemic reduction in circulating autoantibody. Because these chimeras were allowed to reconstitute for approximately 8 weeks before CD21 blockade began, donor B cells had likely already generated and released ssDNA autoantibody into circulation prior to treatment onset; three injections may therefore be insufficient to measurably lower a pre-existing circulating antibody pool, even though GC participation and ASC generation are already suppressed at the cellular level over this same window. Under this interpretation, dual blockade is expected to limit continued autoantibody accumulation rather than deplete antibody already in circulation, a distinction tested directly below with an extended treatment regimen.

**Figure 2.**
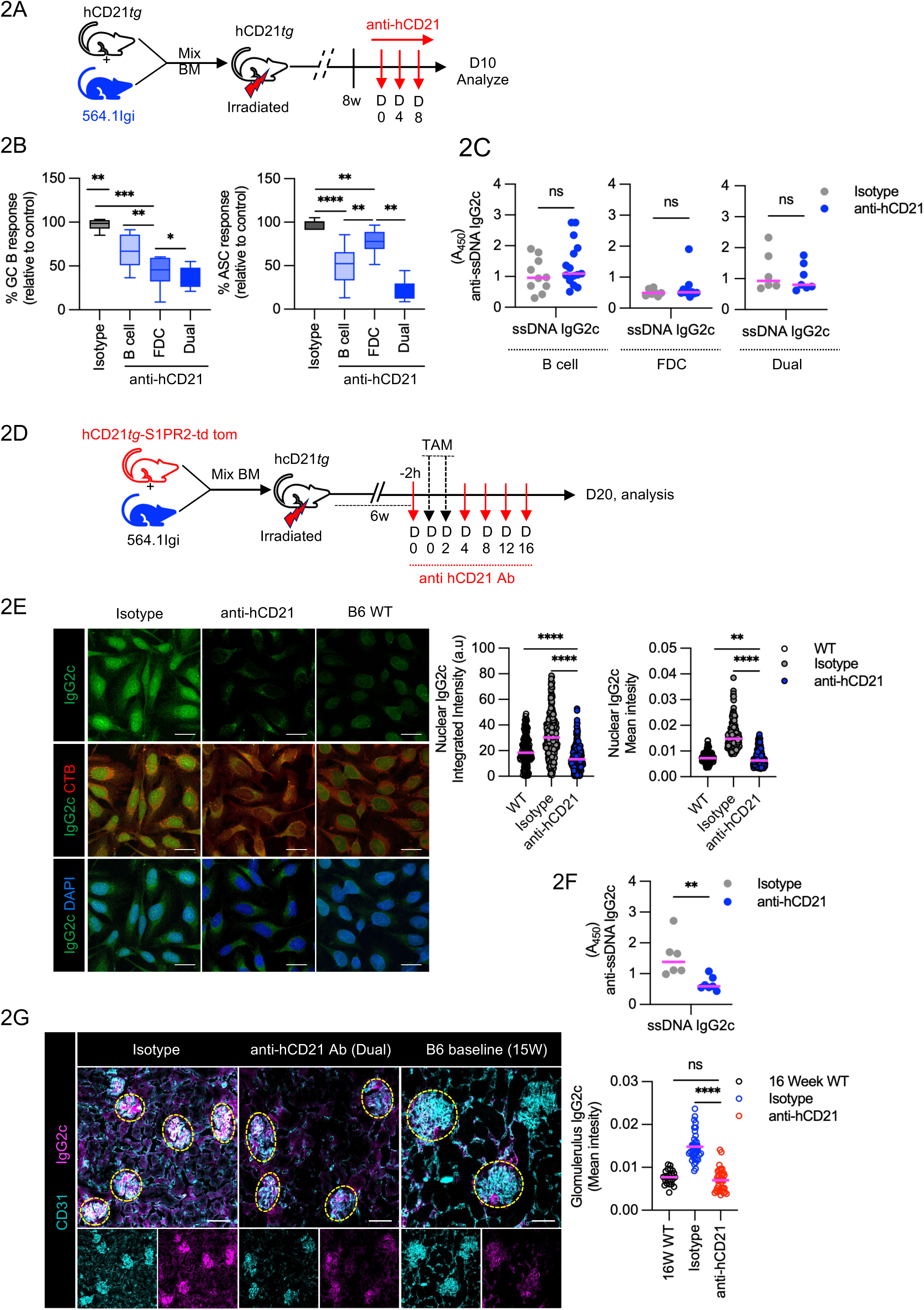
Dual CD21 blockade on B cells and FDCs synergistically suppresses GC B cell and ASC responses, autoantibody production, and renal immune-complex deposition. **A)** Experimental design for dual blockade: hCD21*tg*+564Igi mixed BM chimeras in irradiated hCD21*tg* recipients, three injections of anti-hCD21 antibody (D0, D4, D8), analyzed at D10. **B)** Donor-derived total GC B cell and ASC frequencies (relative to isotype) in single-compartment (B cell or FDC) versus dual blockade. **C)** Serum anti-ssDNA IgG2c autoantibody levels (A_450_) for FDC-, B cell-, and dual-blockade cohorts (isotype versus anti-CD21 Ab). **D)** Experimental design for extended dual blockade: hCD21*tg*-S1PR2-tdTomato+564Igi chimeras in irradiated hCD21*tg* recipients; GC-committed B cells were fate-labeled with TAM administration. Mice received five injections of anti-hCD21 antibody over two weeks and were analyzed at D20. **E)** Representative HEp-2 cell immunofluorescence (IgG2c, cholera toxin B (CTB, cytoplasm), DAPI) and quantification of nuclear IgG2c intensity in WT, isotype, and anti-hCD21 Ab-treated chimeras. Scale bars = 20 μm. **F)** Serum anti-ssDNA IgG2c levels (A_450_), isotype versus anti-hCD21 Ab, following extended dual blockade (5 injections). **G)** Representative kidney immunofluorescence (CD31, IgG2c) in isotype, anti-hCD21 Ab (dual)-treated, and 15-week-old B6 baseline mice, with quantification of glomerular IgG2c deposition. Scale bars = 100 μm. Data represent mean ± SEM of two independent experiments. *p < 0.05, **p < 0.01, ****p < 0.0001 (Mann–Whitney, Prism v10.1). *ns*, not significant. Data combined for female and male mice.

Therefore, next we wanted to test the efficacy of an extended treatment (>3 injections) in reducing systemic accumulation and deposition of donor-derived IgG2c antibodies in organs, specifically in kidneys. We created mixed BM chimeras of 564.1Igi and hCD21*tg*-S1PR2*-tdTomato* mice (hCD21*tg*-*tdTomato*; which carry a Cre-inducible, S1PR2-driven tdTomato fate reporter (S1PR2^cre^ ^Ert2^ ^lox-stop-lox^ td-Tomato; referred to throughout as S1PR2*-tdTomato*) that permanently marks GC-experienced B cells upon tamoxifen (TAM) administration; in irradiated hCD21*tg* recipients to achieve dual blockade (**Fig. 2D**). 6 weeks post-chimera construction activated tdTomato-expressing hCD21*tg* B cells entering GCs were labeled with TAM (oral gavage, see schematic) administration. Mice then received 5 injections (20 μg/mouse i.p.) over 2 weeks and were euthanized on day 20 (D20) post-injection. Control mice received isotype control Ab. As observed before, extending the treatment regimen with the dual-blockade strategy led to a significant and sustained reduction in donor-derived total GC B (GL7^+^CD38^-^) and ASC (CD138^+^GL7^-^) response, (**Figs. S3D and S4A**). Where we observed significant contraction of GC-derived GC B cell (tdTom^+^GL7^+^CD38^-^; **Fig. S3E**); output, GC-derived ASC (tdTom^+^CD138^+^GL7^-^) response did not reach statistical significance (**Fig. 5D**; *data is placed in Fig. 5 for comparison with GC derived tdTom+ ASC response after FDC deletion*), although we observe a trend toward reduced GC derived ASC response. This cellular effect translated into meaningful systemic outcomes, including a substantial decrease in circulating autoreactive antibodies (**Figs. 2E, 2F**); control animals showed a further rise in circulating anti-ssDNA titer over the extended treatment interval, whereas anti-hCD21 Ab-treated animals’ titers remained stable, consistent with dual blockade limiting continued autoantibody accumulation rather than depleting antibody already in circulation. Extended treatment also produced a pronounced reduction in pathogenic IgG2c immune complex deposition in the kidneys, a hallmark of lupus nephritis (**Figs. 2G** **& S4B)**. Thus, sustained dual blockade effectively mitigated both the cellular drivers of disease and its consequential end-organ damage.

Together, these findings demonstrate that dual CD21 blockade produces synergistic suppression of GC B cell and ASC responses, autoreactive antibody accumulation, and renal immune-complex deposition beyond what is achieved by targeting either compartment alone.

### CD21 on B cells is required for initial GC entry and early ASC generation

From these results, however, we couldn’t infer whether reduction in GC B cell (GL7^+^CD138^-^CD38^-^) and CD138^+^ ASC output reflected impaired activation of naïve B cells, impaired GC entry of recently activated B cells or effect of CD21 blockade on B cells already engaged in the GC response (GC-experienced). To determine if CD21 blockade impairs the initial activation and recruitment of recently activated naïve B cells into the autoimmune response, we turned to an adoptive transfer model recently described by van den Broek *et al*. (10) and Zhu *et al*. (39). By doing 2 anti-hCD21 injections, before and after the adoptive B cell transfer, we show that donor B cells can enter an ongoing GC as early as D7 post transfer (**Fig 3A**), and that, blocking CD21 on activated B cells impede their GC entry.

**Figure 3.**
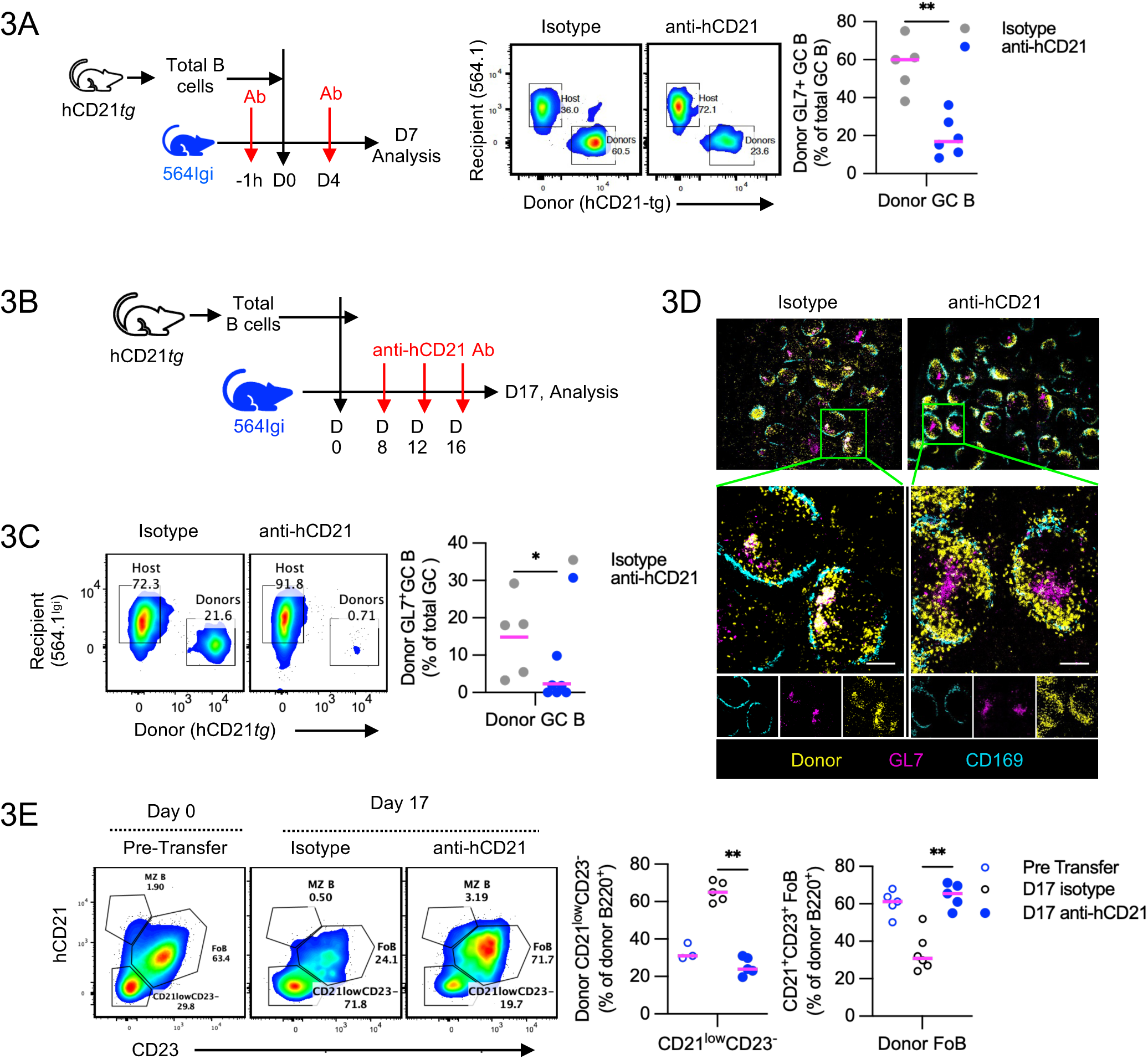
CD21 blockade limits initial activation and GC entry of naïve and recently activated B cells in an adoptive transfer model. **A)** Experimental design for hCD21*tg* total B cells, naïve, transferred into 564Igi recipients, anti-hCD21 antibody given 1 h before and 4 days after transfer, donor GC entry assessed at D7, *left panel*; flow plots and quantification of donor GL7^+^ GC B cells, *right panel*. **B)** Experimental design for a delayed-blockade for addressing the effect of CD21 blockade on GC-derived response; antibody administered beginning D8 (D8, D12, D16), after the initial GC-entry window, analyzed on D17. **C)** Representative flow plots and quantification of donor-derived GC B cells (CD138^-^GL7^+^CD38^-^) at D17. **D)** Representative immunofluorescence showing reduced colocalization of donor B cells (yellow) with GL7^+^ GCs (magenta) in anti-hCD21 Ab-treated versus isotype recipients at D17. Scale bars = 200 μm. **E)** Flow plots (hCD21 versus CD23) at D0 (pre-transfer) and D17, showing follicular (CD21^+^CD23^+^) and activated (CD21^low^CD23^-^) donor B cell subsets, with quantification of each. Data represent mean ± SEM of two independent experiments. *p < 0.05, **p < 0.01 (Mann–Whitney, Prism v10.1).

To further address the effect of CD21 blockade on the capacity of recently activated B cells to drive GC B cells response, and GC dependent-ASCs generation, we adoptively transferred 25 x10^6^ naïve splenic B cells from hCD21*tg* mice intravenously (i.v.) into the 564.1Igi recipients (**Fig 3B**), followed by anti-CD21 antibody treatment. Mice received 3 anti-hCD21 antibody injections day 8 onward, when donor B cells have entered GC. At D17 post-transfer, we found that donor B cells in controls could enter an ongoing GC (acquire GL7 expression) but had reduced capacity to do so with anti-CD21 treatment (**Figs. 3B-3C**). Both donor-derived GC B cells (**Figs. 3C**) and ASC response (**Fig. S5B**) were significantly reduced in antibody-treated animals. Further, the reduced GC B cell response coincided with an impaired ability of donor B cells to enter pre-existing, ongoing GCs in anti-hCD21 Ab-treated animals compared with controls (**Fig. 3D**). Similarly, very few or no donor-derived ASCs were found in red pulp (**Fig. S5C**). Reduced ASC generation coincided with a significant reduction in the production of donor-derived, autoreactive IgG2c antibodies (**Fig. S5E**). These data demonstrate that CD21 co-receptor signaling on naïve B cells is a critical checkpoint for their initial activation, GC recruitment, and early pathogenic ASC differentiation.

Further, in anti-hCD21 Ab-treated mice, CD21 expression was maintained on donor B cells, suggesting limited generation of CD21^low^CD23^-^activated B cells with reduce proliferation potential, reduced CTV dilution and CD138^+^ ASC generation, (**Figs. 3E** **& S5D**) and suggesting that blocking CD21 on naïve B cells significantly limits their initial activation (39). Consistent with the reduced donor-derived GC B cell and ASC responses, CTV analysis at day 17 revealed altered dye dilution patterns following CD21 blockade, suggesting reduced proliferative expansion and/or persistence of highly divided donor B cells (Fig. S5D). Together, these findings support a role for CD21-dependent co-receptor signaling in the expansion and differentiation of recently activated B cells within an autoreactive environment, consistent with previous observations linking CD21 regulation to B cell activation and differentiation (41). This may also contribute to the reduced ASC responses observed in chimeric mice following B cell-associated CD21 blockade.

### CD21 on FDCs Maintains the GC Niche by Mediating Essential B Cell Engagement

While we provide evidence that CD21 blockade on mature B cells limits the GC participation of recently activated B cells, it has little or no effect on B cells engaged in the GC response, as suggested by a modest decrease in GC size with no significant change in GC number per follicle (**Fig. 1E**). While blocking CD21 on FDCs primarily affected GC-specific responses (**Fig. 1D**), the mechanism by which CD21 blockade on FDCs impedes ongoing GC B cell responses is unclear. To address how FDC blockade affects established GCs, we used the S1PR2-tdTomato fate-reporter system (introduced above), to track GC-derived B cell responses, to create mixed BM chimeras of 564.1Igi and S1PR2*-tdTomato* mice in irradiated hCD21*tg* recipients. This allowed specific blockade of hCD21 receptor function on FDCs and tracking of tdTomato^+^ GC B cell generation, GC-derived ASCs, and memory B cell differentiation. At 6 weeks post chimera set-up, activated S1PR2-tdTomato-expressing B cells entering GCs were labeled with TAM (oral gavage, see schematic **Fig. 4A**). Following 4 weeks of TAM treatment, mice received 3 injections of anti-hCD21 Ab (20 μg/mouse i.p.) over 10 days and were euthanized at D10 post-blockade. Control mice received isotype Ab.

**Figure 4.**
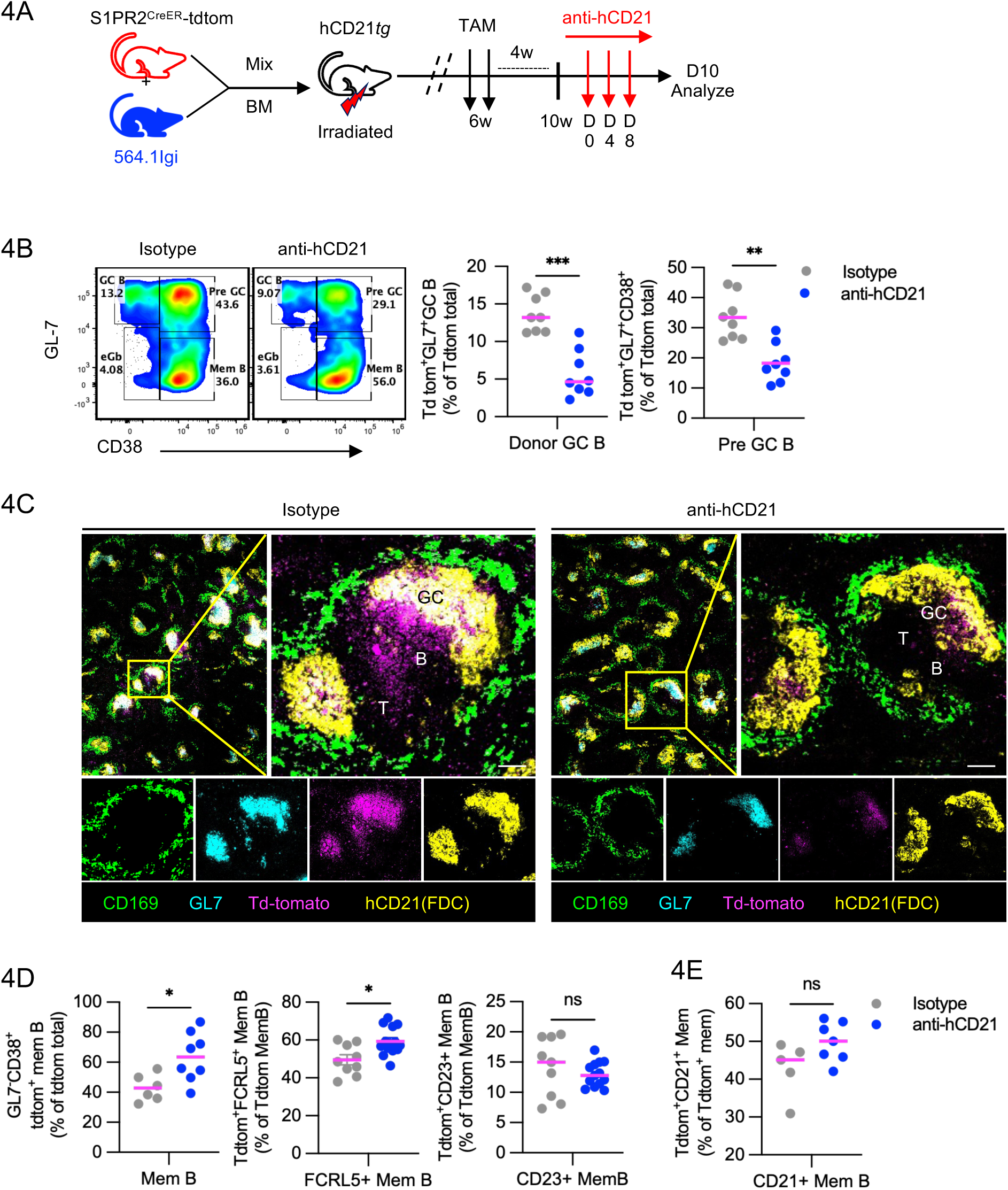
FDC-specific CD21 blockade disrupts GC B cell–FDC engagement and destabilizes established GCs. **A)** Experimental design to address the effect of GC niche disruption on GC derived response outcome; 564.1Igi+S1PR2-tdTomato chimeras in irradiated hCD21*tg* recipients; GC-committed B cells were fate-mapped with TAM from 6 weeks. Mice received anti-hCD21 antibody from 10 weeks (D0, D4, D8) after memory is established and were euthanized at D10 to analyze GC derived responses. **B)** Flow plots and quantification of donor-derived (Td-tomato^+^) GC B cells (GL7^+^CD38^-^) and pre-GC B cells (GL7^+^CD38^+^). **C)** Representative immunofluorescence (CD169, GL7, Td-tomato, hCD21(FDC)) showing GC B cell-FDC colocalization in isotype versus anti-hCD21 Ab-treated chimeras. **D)** Flow plots and quantification of donor-derived (Td-tomato^+^) memory B total (CD38^+^GL7^-^) and memory subsets (FCRL5+, CD23+ memory subset). **E)** Quantification of GC-derived (Td-tomato^+^) CD21^+^ memory B cell (GL7^-^CD21^+^CD38^+^). Data represent mean ± SEM of two independent experiments. *p < 0.05, **p < 0.01 (Mann–Whitney, Prism v10.1). *ns*, not significant.

**Figure 5.**
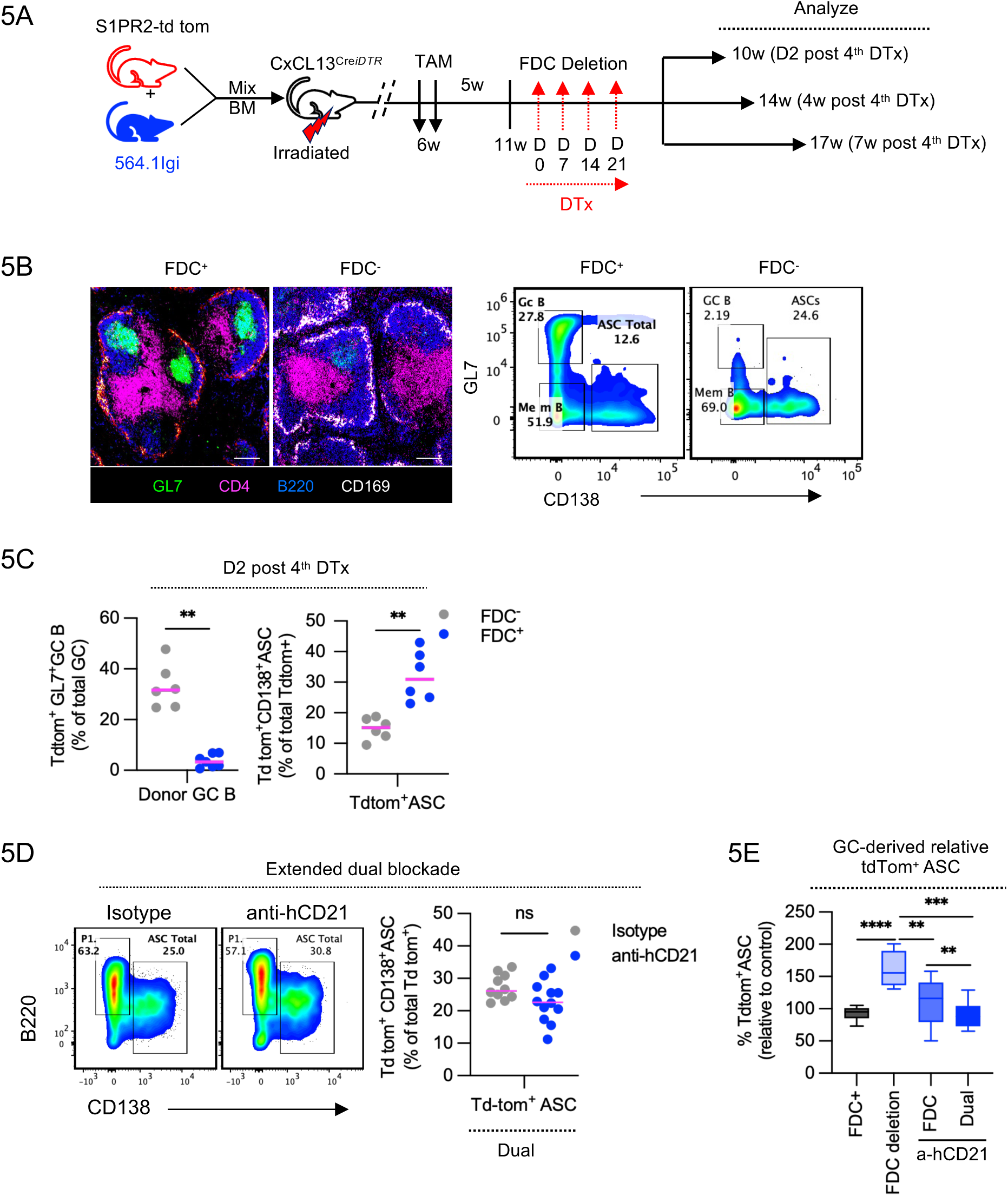
Disruption of the GC niche by FDC deletion exacerbates GC-derived ASC differentiation, an effect contained by dual CD21 blockade. **A)** Experimental design to address the effect of FDC deletion on GC derived response outcome; S1PR2-tdTomato+564.1Igi chimeras in irradiated CXCL13^CreDTR^ recipients; GC-committed B cells were fate-labeled with TAM at 6 weeks. Mice received diphtheria toxin (DTx, four doses) from 11 weeks to induce transient FDC deletion and were analyzed at 10, 14, and 17 weeks (D2, 4 weeks, and 7 weeks after the 4th DTx dose). **B)** Representative immunofluorescence (GL7, CD4, B220, CD169) and flow plots (GL7 versus CD138) comparing FDC-intact (FDC^+^) and FDC-deleted (FDC^-^) mice, with quantification of GC-derived (Td-tomato^+^) ASCs at D2 post-4th DTx dose. **C)** GC-derived GC B (GL7^+^CD38^-^) and ASC (CD138^+^GL7^-^) frequency following FDC deletion versus **D)** pharmacological dual CD21 blockade showing GC derived ASC (tdTom^+^CD138^+^GL7^-^) frequencies. **E)** GC-derived ASC frequency (relative to isotype) across FDC-deletion, FDC-blockade, and dual-blockade conditions. Data represent mean ± SEM of two independent experiments. *p < 0.05, **p < 0.01, ***p < 0.001, ****p < 0.0001 (Mann–Whitney, Prism v10.1). *ns*, not significant.

Consistent with our earlier observations (**Fig. 1**), FDC specific CD21 blockade significantly reduced donor-derived total GC B (GL7^+^CD38^-^) but not ASC (CD138^+^GL7^-^) response, (**Fig. S6A)** confirming the effect of FDC specific blockade in reducing GC response. Consequently, FDC-specific CD21 blockade in chimeric mice critically reduced the number of ongoing GC-derived donor B cell (tdTomato^+^ GL7^+^CD38^-^) and pre-GC B cells (tdTomato^+^GL7^+^CD38^+^) response (**Fig. 4B**) and significantly abrogated their physical colocalization with FDCs (CD21/35^+^) within the niche (**Figs. 4C** **& S6C**), further demonstrating that blockade of CD21 on FDCs significantly ablated contact between GC B cells and FDCs. This disruption of the GC microenvironment coincided with significantly more memory B cells (MBCs) accumulation, (**Fig. 4D, *left panel***). Further characterization of the GC-derived memory B cell compartment revealed a significant increase in the FCRL5^+^ memory B cell subset following FDC-associated CD21 blockade, whereas the CD23^+^ memory B cell subset remained relatively unchanged (**Fig. 4D**). Previous work by Castrillon et al. (40) identified FCRL5^+^ and CD23^+^ as distinct GC-derived memory B cell populations with different transcriptional profiles and splenic localization. Given the reported propensity of FCRL5^+^ memory B cells to rapidly differentiate into antibody-secreting cells (ASCs) upon recall, their accumulation following FDC-associated CD21 blockade is particularly intriguing. It will be interesting to determine whether these accumulated FCRL5^+^ memory B cells contribute to subsequent ASC responses and whether CD23^+^ memory B cells preferentially retain the capacity for GC re-entry. These observations raise the possibility that disruption of the FDC-supported GC niche influences not only GC maintenance but also the composition and functional potential of the resulting memory B cell compartment.

Over 50% of these GC-derived MBCs retained CD21 expression (**Fig. 4E**), suggesting a heightened potential for ASC differentiation upon recall (antigen re-encounter), a property directly tested in later experiments (**Fig. 6C**). Unlike dual blockade where GC-derived ASC (tdTom^+^CD138^+^GL7^-^) response remains unchanged in Ab treated vs control animals (**Fig. 5D**), mice with FDC-specific blockade showed a trend towards increased accumulation of GC-derived ASC (tdTom^+^CD138^+^GL7^-^), **Fig. S6B**.

**Figure 6.**
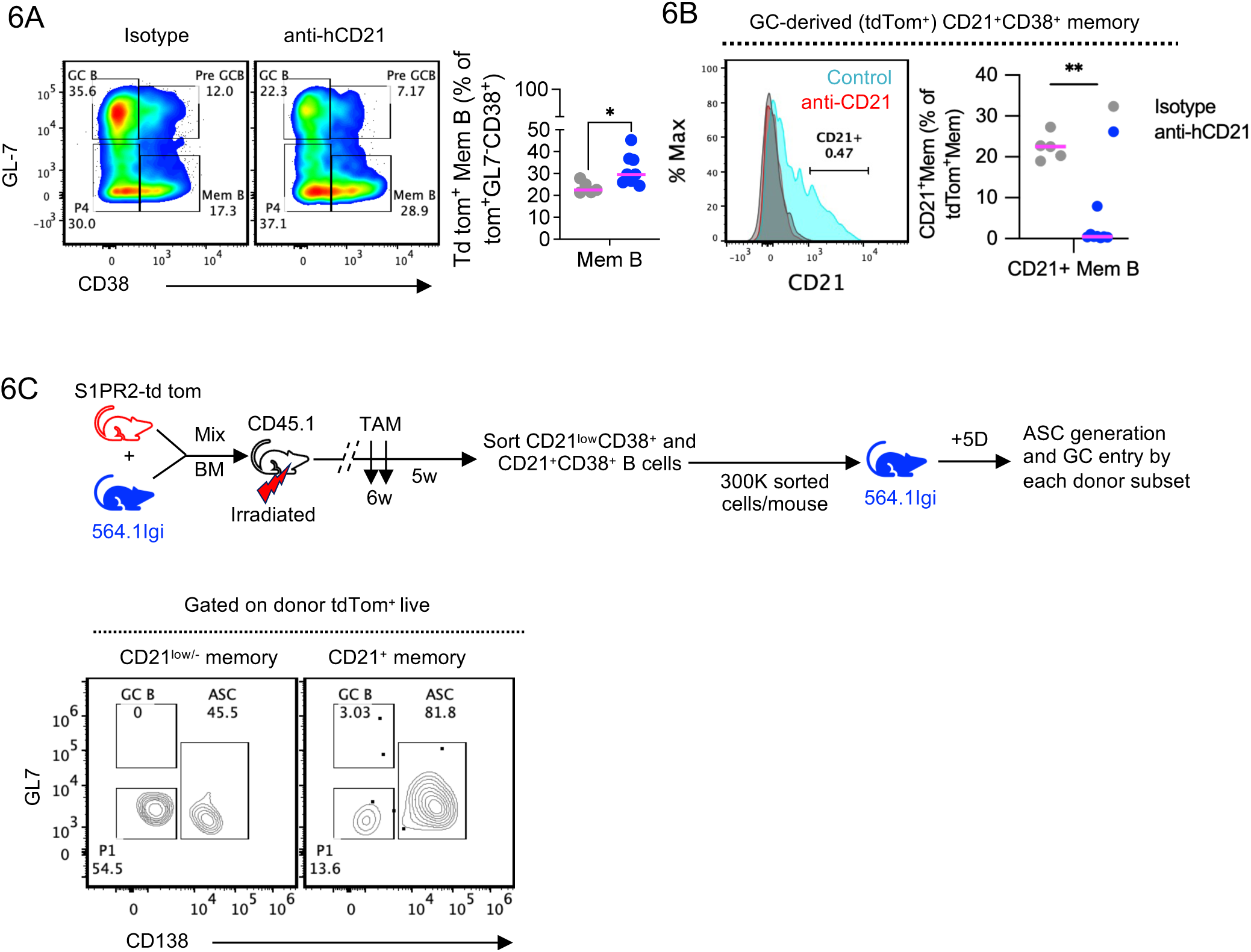
Dual CD21 blockade promotes accumulation of a GC-derived CD21^low/-^CD38^+^ memory B cell subset with reduced ASC-generating capacity. A) Representative flow plots (GL7 vs CD38) and quantification showing GC-derived (tdTomato^+^) memory B cell (GL7^-^CD38^+^) frequency, isotype versus anti-hCD21 Ab. B) Representative CD21 histogram and quantification of the proportion of GC-derived memory B cells retaining CD21 expression (CD21^+^CD38^+^) in dual CD21 blockade condition. C) Experimental set-up for functional recall: CD21^low^CD38^+^ and CD21^+^CD38^+^ memory B cells sorted from S1PR2-tdTomato/564.1Igi chimeras and transferred (300,000 cells/mouse) into naïve 564.1Igi recipients; ASC generation assessed 5 days later. Flow plots and quantification of donor-derived CD138^+^ ASCs from each subset. Data represent mean ± SEM of two independent experiments. *p < 0.05, **p < 0.01 (Mann–Whitney, Prism v10.1).

### Complete FDC deletion alone is insufficient for maximal therapeutic suppression

The accumulation of MBCs and ASCs after the collapse of established GCs is consistent with GC-engaged B cells being diverted toward the ASC differentiation pathway following loss of niche support. To test whether GC niche disruption inherently exacerbates ASC output, we performed selective FDC deletion experiments. We created mixed BM chimeras of 564.1Igi and S1PR2*-tdTomato* in irradiated CxCl13^Cre^DTR recipients. The BM allowed us to track tdTomato^+^ B cells in an autoreactive environment, while the CxCl13^Cre^DTR recipient allowed us to transiently deplete FDCs using diphtheria toxin (DTx) (**Fig. 5A**). 6 weeks post-chimera construction activated S1PR2-tdTomato-expressing B cells entering GCs were labeled with TAM (oral gavage, see schematic) administration. 5 weeks post TAM treatment, when GC derived-Tdtom^+^ memory B compartment is established, mice received 4 injections of diphtheria toxin (DTx; 2 μg/ml i.p.) and were euthanized at the indicated times. FDC deletion led to complete GC dissolution and a significant increase in the number of GC-derived tdTomato^+^ ASCs (**Figs. 5B, 5C**), compared to pharmacological blockade (**Figs 5D-5E)**. This confirms that isolated disruption of the GC niche, while halting GC activity, has the unintended consequence of accelerating the conversion of GC-experienced B cells into pathogenic ASCs. Critically, this compensatory ASC response was not observed during dual CD21 blockade on both B cells and FDCs (**Fig 5D**). In this coordinated intervention, collapse of the GC niche (via FDC targeting) was coupled with direct inhibition of B cell differentiation capacity (via B cell targeting), producing a synergistic reduction of GC B cells and total/GC-derived ASCs, with no compensatory ASC increase. Thus, dual blockade uniquely contains the pathogenic ASC output unchecked by isolated GC niche disruption, demonstrating that coordinated targeting of both cellular compartments is required to comprehensively suppress the autoimmune effector response.

### Dual CD21 blockade remodels the autoreactive memory compartment

Because memory B cells (MBCs) generated through autoreactive GC reactions can drive rapid, pathogenic antibody production upon recall (3,4,42–45), we next characterized the memory compartment generated following CD21 blockade. Dual CD21 blockade (done as in **Fig. 2D**), unlike FDC blockade alone, promoted the accumulation of a CD21^low/-^CD38^+^ MBC subset (**Figs. 6A-6B, S7B**). Total GC-derived (Td-tomato+) memory B cell frequency was significantly increased following both FDC and dual CD21 blockade (**Fig. S7A**). To test whether this CD21^low/-^memory subset is intrinsically less capable of generating ASCs, independent of ongoing CD21 blockade, we generated hCD21-S1PR2-*tdTomato*+564.1Igi mixed BM chimeras in irradiated B6.1 (CD45.1) hosts, without anti-CD21 antibody treatment, and sorted donor-derived (Td-tomato^+^) CD21^+^CD38^+^ and CD21^low/-^CD38^+^ GC-experienced memory B cells by congenic marker expression. Each sorted subset was then adoptively transferred separately into naïve 564.1Igi recipients to ask whether CD21 expression status at the time of transfer intrinsically predicts ASC-generating capacity in an autoreactive environment (**Fig. 6C**).

Functional assessment showed that the CD21^low^CD38^+^ memory subset had a reduced capacity to differentiate into ASCs when transferred to the 564igi host (**Fig. 6C**). Instead, we found that the CD21^+^CD38^+^ memory B subset generated more ASCs *in vivo*. This finding provides a mechanistic basis for the reduced autoreactive antibody output observed with dual blockade: the shift toward a CD21^low/-^CD38^+^ memory compartment is not merely a passive consequence of blockade but reflects accumulation of a population that is intrinsically poor at generating pathogenic ASCs, directly linking the memory remodeling induced by dual CD21 blockade to its suppression of autoreactive antibody production.

## Discussion

Persistent autoreactive germinal center (GC) responses represent a major therapeutic challenge in systemic lupus erythematosus (SLE) because they provide an evolving reservoir in which autoreactive B cell clones can expand, diversify, recruit additional B cells, and continuously generate pathogenic humoral effector populations (7–10, 40). In this study, we identify CD21 as a regulatory node operating at two mechanistically distinct cellular compartments: B cells, where CD21 amplifies activation through the BCR co-receptor complex, and follicular dendritic cells (FDCs), where CD21 supports antigen retention and maintenance of the GC niche. Our findings demonstrate that maximal suppression of lupus-associated humoral immunity requires coordinated targeting of these two CD21-dependent processes, thereby simultaneously limiting autoreactive B cell recruitment, destabilizing established GC responses, suppressing ASC generation, and altering pathogenic memory recall.

A central finding of this study is the mechanistic dichotomy between B cell-intrinsic and FDC-associated CD21 function. Selective blockade of CD21 on B cells effectively impaired activation-dependent recruitment of naïve autoreactive B cells into ongoing GC responses and reduced ASC differentiation. These findings are consistent with the established ability of the CD19-CD21-CD81 co-receptor complex to markedly lower the threshold for BCR activation following recognition of complement-opsonized antigen (27–31). They also extend the recent findings of Zhu et al., in which continuous CD21-complement engagement preceded proliferation of naïve autoreactive B cells and functional CD21 blockade suppressed progression toward the extrafollicular ASC program (39). Together, these observations suggest that CD21 does not simply enhance the magnitude of an established B cell response but can operate at an early checkpoint controlling whether subthreshold or low-affinity autoreactive BCR engagement progresses toward productive activation.

This mechanism may be particularly important in lupus because autoreactive GCs are not closed structures populated exclusively by founder clones. Rather, established spontaneous GCs can continuously recruit naïve B cells, which subsequently clonally expand and diversify the autoimmune repertoire (10). Restricting CD21-dependent activation may therefore reduce the continual influx of new autoreactive clones into both GC and extrafollicular pathways. Such a mechanism provides a potential explanation for why B cell-restricted CD21 blockade strongly suppresses newly generated ASC responses while exerting a more limited effect on GCs that are already established.

In contrast, selective blockade of CD21 on FDCs produced a mechanistically distinct phenotype characterized by profound disruption of GC architecture and collapse of the GC niche.

Importantly, however, isolated disruption of the GC niche was insufficient to extinguish pathogenic humoral immunity. Instead, selective FDC blockade and GC collapse were accompanied by increased accumulation of GC-derived ASCs, suggesting that interruption of the GC program can redirect GC-engaged B cells toward terminal differentiation. This finding is strongly supported by Voss et al., who genetically blocked the GC pathway through loss of BCL6 and demonstrated that GC ablation failed to prevent autoimmune progression in both TLR7-driven and 564Igi lupus models. Rather, GC blockade accelerated plasma-cell differentiation and produced a disproportionate contribution of GC-blocked B cells to the plasma-cell compartment (46). Thus, removing the GC niche does not necessarily eliminate autoreactive B cells that have already received sufficient activation signals; instead, it can alter their differentiation trajectory. Together, these findings argue that intervening solely on the GC-dependent pathway is unlikely to be sufficient to limit autoantibody production and curtail SLE pathogenesis.

This concept is particularly important because extrafollicular and GC pathways should not be viewed as mutually exclusive sources of pathogenic autoantibodies. Human and experimental SLE can generate substantial ASC responses through extrafollicular pathways, and disease flares are associated with large expansions of activated naïve-derived and DN2-like effector B cell populations capable of rapid plasma-cell differentiation (47–49). TLR7-driven CD11c^+^ B cell populations likewise accumulate in murine autoimmunity and provide an additional mechanistic connection between innate nucleic-acid sensing and extrafollicular B cell differentiation (50).

The recent demonstration that CD21 primes autoreactive extrafollicular differentiation (39), together with our observation that isolated GC disruption can increase ASC output, therefore suggests that therapeutic targeting of only one differentiation pathway may permit functional escape through the other.

The synergistic efficacy of bicellular CD21 blockade likely reflects simultaneous suppression of these complementary pathogenic mechanisms. Whereas B cell CD21 blockade restricts activation and recruitment of newly responding autoreactive cells, disruption of FDC-associated CD21 removes the antigen-retaining architecture required to sustain established GC reactions.

By targeting these processes concurrently, dual blockade prevented the compensatory ASC accumulation observed following isolated FDC intervention and produced robust suppression of GC responses, autoreactive ASC generation, circulating autoantibodies, and renal immune-complex deposition. These findings support a broader therapeutic principle: effective interruption of chronic humoral autoimmunity may require simultaneous targeting of the pathogenic lymphocyte and the tissue niche that sustains its continued evolution.

An additional and particularly important finding of this study is the impact of coordinated CD21 blockade on the autoreactive memory compartment. Prolonged dual CD21 blockade promoted accumulation of a CD21^low/−^CD38^+^ memory B cell subset with markedly reduced ASC recall potential, whereas isolated FDC blockade enriched CD21^+^CD38^+^ GC-derived memory populations that retained strong ASC differentiation capacity. Memory B cells are highly heterogeneous and can be developmentally and functionally programmed toward markedly different secondary-response fates (45). Early studies demonstrated that distinct memory populations preferentially regenerate either plasma cells or secondary germinal centers upon recall (51). CD80 and PD-L2 expression subsequently defined functionally distinct murine memory B cell populations with differential propensity for rapid ASC differentiation versus GC re-entry (52). GC output itself is temporally regulated, with memory B cells preferentially generated earlier and plasma cells increasingly produced during later stages of the GC response (53). Collectively, these findings established that memory B cells do not constitute a uniform reservoir but instead comprise populations preconfigured for divergent recall responses (44,45,51–53). Recent lineage and epigenomic studies have further strengthened this concept. Memory B cell subsets can arise from distinct developmental histories that imprint different chromatin and transcriptional states and subsequently constrain their capacity to re-enter GCs or differentiate into plasmablasts (54). More broadly, memory B cells exist along a functional continuum in which subset identity, activation history, and differentiation state influence their probability of generating plasma-cell versus GC responses following secondary stimulation (55). These observations provide an important framework for interpreting the memory remodeling induced by CD21 blockade. Our findings suggest that the cellular site of CD21 intervention influences not only the magnitude of ongoing autoreactive GC responses but also the subsequent developmental state and pathogenic potential of the memory compartment. Thus, coordinated CD21 blockade may provide durable benefit not simply by suppressing the active autoimmune response but by changing the quality of the B cell populations that survive treatment.

Importantly, however, the CD21^low/−^CD38^+^ population generated in our experimental system should not be equated broadly with CD21^low^ populations described in human SLE. Human CD21^low/-^atypical and DN2-like B cell populations are heterogeneous, and several subsets display enhanced TLR7 responsiveness, T-bet expression, inflammatory transcriptional programs, and strong plasma-cell differentiation potential (49,56). Human autoreactive B cell states are similarly shaped by multiple tolerance checkpoints and activation pathways that cannot be defined by CD21 expression alone (57). The reduced recall capacity observed here should therefore be interpreted as a functional property of the specific CD21^low/−^CD38^+^ population generated following dual blockade rather than as a general property of CD21^low^ B cells.

Transcriptional, epigenetic, and clonal lineage analyses will be important for defining how this population relates to previously characterized memory, atypical-memory, and pre-ASC states.

These findings have important translational implications in the context of current lupus therapies. Rather than globally eliminating the B cell lineage, CD21 blockade targets molecular interactions required to initiate and sustain complement-dependent autoreactive humoral responses. Simultaneous interference of B cell co-receptor activation and FDC-supported antigen persistence may therefore dismantle a pathogenic immune circuit while preserving B cells that are not actively dependent on this pathway. Whether such an approach preserves protective immunity more effectively than broad B cell depletion remains to be determined and will require direct assessment of primary responses to foreign antigens, vaccine responses, affinity maturation, and immunological memory.

Several additional questions remain. Although CD21 blockade did not cause overt B cell depletion in our models, CD21 contributes to physiological humoral responses, and prolonged blockade could potentially alter affinity maturation or responses to limiting antigen (27–36). Translation to human SLE must also account for differences in CR1/CR2 expression and function between mouse and human immune systems. Furthermore, the durability, transcriptional identity, developmental origin, and clonal relationships of the CD21^low/−^CD38^+^ memory population remain to be defined. Additionally, because the functional recall assay transferred sorted memory subsets exclusively into autoreactive 564.1Igi hosts, whether placing these subsets into a non-autoimmune host would alter the degree of autoreactive ASC generation remains untested, an important caveat for distinguishing pathogenic potential intrinsic to each subset from that conferred by the recipient’s autoreactive environment. Determining whether analogous memory remodeling occurs in human B cells after CD21 inhibition will be particularly important for therapeutic development.

In summary, our study establishes CD21 as a dual-node regulator of pathogenic humoral immunity. B cell–associated CD21 controls activation-dependent recruitment and ASC differentiation, whereas FDC-associated CD21 maintains the antigen-rich GC niche required for persistence of established autoreactive responses. Disrupting only the FDC/GC node permits compensatory ASC output, whereas coordinated blockade of both compartments suppresses GC and extrafollicular escape pathways and reshapes subsequent memory recall. These findings identify the CD21-mediated B cell-FDC axis as a therapeutically actionable circuit in lupus and provide a broader conceptual framework in which effective treatment of chronic autoimmune responses may require simultaneous targeting of pathogenic immune cells and the tissue niches that sustain them.

## Materials and methods

### Mice and Ethics statement

C57BL/6J wild-type (B6 WT), B6.SJL (CD45.1) and tdTomato (stock number 007914) reporter mice were obtained from Jackson Laboratories (USA). mCD21 knockouts (mCD21^-/-^) were bred and maintained in-house. Human CD21-transgenic (expressing the transgenic hCD21 receptor; hCD21*tg*; hCD21^+^mCD21^-/-^) mice were provided by Dr. Michael V. Holers (University of Colorado School of Medicine, CO, USA). ROSA26iDTR mice (iDTR;007900) were purchased from Jakson. Cxcl13-Cre/Tdtomato [Tg(Cxcl13-cre,-tdTomato)719Biat, EMMA repository catalog no. EM:13610] were obtained from the European Mouse Mutant Archive (58, 59). To perform conditional follicular dendritic cell (FDC) ablation, Cxcl13-Cre/Tdtomato mice were crossed with iDTR, replicating previously described targeting strategy (60). 564Igi mice (Berland et al., 2006) were originally provided by Theresa Imanishi-Kari (Tufts University) and were maintained in-house. S1pr2-CreERT2 BAC-transgenic mice (Shinnakasu et al., 2016) were generously provided by T. Kurosaki (RIKEN-Yokohama). 564.1Igi mice were generated by crossing 564Igi and B6.SJL (CD45.1) mice. S1PR2-Cre^ERT2^ ^lox-stop-lox^-Td-Tom (S1PR2-*tdTom*) fate reporter mice were created by crossing S1pr2-CreERT2 BAC-transgenic mice with a tdTomato fate reporter. hCD21^+^mCD21^-/-^mice were generated by crossing hCD21*tg* mice with mCD21^-/-^mice and were maintained in-house. All mice were bred and maintained in the AAALAC-accredited facility at Harvard Medical School. Mice were specific pathogen-free and maintained under a 12-hr light/dark cycle with a standard chow diet. Both male and female mice were used. All animal experiments were conducted in accordance with the guidelines of the Laboratory Animal Center of the National Institutes of Health. The Institutional Animal Care and Use Committee of Harvard Medical School approved all animal protocols (protocol number IS00000111). All mice were genotyped by ddPCR/PCR using the primers listed in **Table 1**.

### Antibodies, staining reagents and immune cell kits

All the antibodies used for immunofluorescence microscopy or flow cytometry analysis are listed in **Table 2**. For total B cell negative enrichment, MojoSort kits (cat# 480088) were purchased from Biolegend USA. Viability dye for live/dead (Fixable Viability Dye eFluor 780, cat# 65-0865-18) staining was purchased from eBioscience, USA. Monoclonal hCD21 blocking antibody (clone 658) was generated in collaboration with Pfizer Inc., USA (61).

### Receptor occupancy (RO) assay

To quantify in vivo CD21 receptor occupancy, blood or splenic B cells from anti-hCD21 antibody-treated and isotype control mice were stained in parallel with two directly fluorochrome-conjugated anti-hCD21 antibodies recognizing distinct, non-overlapping epitopes:

i. the same blocking clone (658), conjugated to Alexa Fluor-647, which competes with previously administered unlabeled blocking antibody for binding and therefore reports the frequency of unoccupied hCD21 receptors; and (ii) the non-blocking clone Bu32, which binds irrespective of blockade status and therefore reports total surface hCD21 expression as a normalization control. Percent receptor blockade was calculated from the reduction in clone-658 (AF647) staining frequency or mean fluorescence intensity in antibody-treated animals relative to isotype-treated controls, normalized to total receptor expression detected by clone Bu32 staining in the same sample.

### Bone marrow (BM) chimera construction

BM chimeras were prepared using marrow from 564Igi mice and different WT donor mice as previously described (9). Briefly, BM recipient mice were lethally irradiated at 1100 rad and kept on antibiotics (sulfamethoxazole) through drinking water for 10 days after irradiation. For some experiments, mice received 2 doses of 550 rad/cycle, 4 hour apart. For BM construction, femurs and tibias from donor mice were harvested, and marrow was extracted from bones by crushing them using a mortar and pestle in BM cell transfer buffer (HBSS supplemented with 10 mM N-2-hydroxyethylpiperazine-N-2-ethanesulfonic acid (HEPES), 1 mM Ethylenediaminetetraacetic acid (EDTA), and 2% heat-inactivated fetal bovine serum). After RBC lysis (as described below), the cell pellet was resuspended in HBSS buffer and passed through a 70-μm sterile filter. All autoimmune chimeras were prepared at a 2:1 ratio for WT:564Igi donors. Irradiated hosts received a total of 21 × 10^6^ donor mix BM cells by single retro-orbital injection in 100 μl HBSS buffer using a 1 mL insulin syringe and a 29G needle (Becton Dickinson), approximately 8 h post-irradiation. Compartment-specific CD21 blockade exploited the fact that FDCs are radioresistant stromal cells that are not replaced by donor bone marrow, whereas the B cell compartment is fully reconstituted from donor marrow; placing the hCD21^+^mCD21^-/-^(hCD21*tg*) genotype in the donor versus the recipient of a given chimera therefore restricted human CD21 expression, and thus anti-hCD21 antibody binding, to either the B cell compartment or the FDC compartment, respectively.

### FDC depletion

To transiently delete follicular dendritic cells (FDCs), Cxcl13-Cre mice were crossed with iDTR mice to generate Cxcl13Cre-DTR mice, in which Cre-mediated DTR expression is constitutive in Cxcl13-expressing stromal cells, including FDCs. Both transgenes were genotyped by PCR using the primers in Table 1. Lethally irradiated Cxcl13Cre-DTR recipients were reconstituted with a mixture of S1pr2-CreERT2-tdTomato and 564.1Igi bone marrow, as described above.

Because FDCs are radioresistant and remain of host origin, DTR expression was restricted to the recipient stromal compartment, whereas the B cell compartment derived from donor marrow. Six weeks after reconstitution, mice received two oral gavages of 200 μL TAM (50 mg/mL in sunflower seed oil) to label S1PR2-expressing GC B cells and their progeny with tdTomato.

After memory B cells had formed (5 weeks after TAM), mice received four intraperitoneal injections of diphtheria toxin (DTx; 100 μl of 2 µg/mL stock) to transiently deplete FDCs and were euthanized at the indicated times. GC B cells (GL7^+^CD38^−^), memory B cells (CD38^+^GL7^−^) and ASCs (CD138^+^), both total and tdTomato^+^, were quantified by flow cytometry littermate CxCl13Cre^-/-^iDTR^+/+^ mice were used as controls.

### Adoptive B cell transfer

For adoptive transfer, 564.1Igi mice intravenously (i.v.) by retro-orbital injection (100 μL, single eye, 0.5 mL insulin syringe with 29G needle) received MACS-enriched splenic B cells (25 × 106 cells/mouse) isolated from hCD21tg mice. Briefly, donor splenocytes from hCD21tg mice were harvested and minced to prepare single-cell suspensions, and RBCs were removed using ammonium-chloride-potassium (ACK) lysis buffer. The cells were then washed with PBS and passed through a 70 μm-pore cell strainer. B cells were negatively enriched using the MojoSort Mouse Pan B Cell Isolation Kit (Biolegend, cat# 480088) according to the manufacturer’s instructions. Briefly, splenocytes (100 × 10^6^ cells/mL in MACS buffer) were incubated for 15 min on ice with the biotinylated antibody cocktail and then for 15 min on ice with streptavidin magnetic beads, and applied to an LS column (Miltenyi Biotec, cat# 130-042-401) placed in a magnetic field. The unlabeled flow-through fraction was collected, counted, and resuspended in HBSS supplemented with 10 mM HEPES, 1 mM EDTA and 2% FBS for injection. Purity of B cells (B220^+^CD3e^-^cells) was confirmed using BD FACS Canto (BD Biosciences, San Jose, CA). The purity of these sorted cells was >96%. Indicated doses of Anti-hCD21 blocking antibody (clone 658) or human IgG1/κ isotype control (BD Biosciences, cat# 569605) was administered at indicated times by intraperitoneal route.

### Cell Trace Violet labeling for in vivo B-cell proliferation analysis

To assess the effect of CD21 blockade on the proliferation of recently activated B cells, naïve splenic B cells were enriched from hCD21tg donor mice and labeled with Cell Trace Violet (CTV; Thermo Fisher Scientific) according to the manufacturer’s recommended protocol.

Briefly, a 5 mM CTV stock solution was prepared by reconstituting the dye in 20 μL of dimethyl sulfoxide (DMSO). A 5 μM working solution was prepared by diluting 10 μL of CTV stock in 10 mL of phosphate-buffered saline (PBS). Enriched B cells were washed, resuspended in PBS, and incubated with the CTV working solution for 10 min at 37°C, with gentle mixing by tube inversion approximately every 5 min. Following incubation, cells were placed on ice, and staining was quenched by adding 3–4 volumes of RPMI medium supplemented with 10% fetal bovine serum (FBS). Labeled cells were centrifuged at 1,400 rpm for 5 min at 4°C, resuspended in RPMI containing 10% FBS, and counted. Subsequently, 25 × 10⁶ CTV-labeled naïve B cells were collected, washed, and resuspended in 100 μL of HBSS supplemented with 10 mM HEPES, 1 mM EDTA, and 2% FBS. Cells were adoptively transferred intravenously into 564.1Igi recipient mice (day 0). Following adoptive transfer, recipient mice received three intraperitoneal injections of anti-hCD21 blocking antibody or isotype control according to the treatment schedule illustrated in **Fig. 3B**. Mice were euthanized on day 17 post-transfer, and spleens were collected for flow cytometric analysis. Donor-derived B cells were identified using congenic markers, and their proliferation was assessed by CTV fluorescence dilution. CD138 expression was analyzed alongside CTV fluorescence to characterize the proliferative response of donor-derived B cells following CD21 blockade (**Fig. S5D**).

### Sorted memory B cell adoptive transfer and functional recall assay

Fate mapped GC-derived (Td-tomato^+^) memory B (MBC) cells (CD38^+^GL7^-^) were generated in S1PR2-tdTomato/564.1Igi mixed BM chimeras established in irradiated WT (CD45.1or hCD21*tg*) recipients, allowing donor-derived (Td-tomato+) memory subsets to be unambiguously identified and sorted irrespective of host background. These cells were further subdivided into CD21^low/-^and CD21^+^ subsets by fluorescence-activated cell sorting. Sorted populations were washed, counted, and resuspended in HBSS, and 300,000 cells of each subset were adoptively transferred i.v., without further anti-CD21 antibody treatment, into naïve

564.1Igi recipient mice to test the cell-intrinsic capacity of each memory subset to differentiate into ASCs upon reintroduction into an autoreactive environment. Recipients were euthanized 5 days post-transfer, and donor-derived (Tdtomato^+^) ASC (CD138^+^) and GC B cell (GL7^+^CD38^-^) generation were assessed by flow cytometry as described above.

### Tracking GC-experienced and GC-derived B cells, tissue collection and processing

For tracking GC-experienced B cells, BM from S1PR2-tdTomato and hCD21*tg*-S1PR2-tdTomato fate reporter mice was used to generate a mixed BM chimera with 564Igi donors into irradiated hosts. GC fate-committed B cells were tracked by inducing tdTomato expression in S1PR2-expressing B cells. Cre-inducible tdTomato expression in S1PR2^CreER^tdTomato donor B cells was achieved by TAM treatment of BM recipient mice. TAM solution was prepared by dissolving TAM powder in sunflower seed oil (Sigma) O/N at 37 °C in the dark. Mice received 2 doses of 200 μl of 50 mg/mL TAM solution orally (gavage), as described in the results. For tissue collection, mice were euthanized at indicated time points by cervical dislocation under isoflurane-induced anesthesia. Spleens, lymph nodes and kidneys were harvested and processed for adoptive transfer, immunofluorescence (IF) microscopy and flow cytometry analysis (see below). Bone marrow (BM) was collected for the construction of chimeric mouse models (see below). For preparing single-cell suspensions from spleen and BM, the cell pellet obtained after tissue processing was incubated in RBC lysis buffer (155 mM NH_4_Cl, 10 mM NaHCO_3_, 0.1 M EDTA) for 5 min at room temperature (RT) to lyse RBCs. Cells were spun at 1500 rpm for 5 min. Serum was collected for quantifying donor-derived autoantibody responses by antigen-specific ELISA and Hep2 staining (see below). For IF staining, half-spleens were flash-frozen on dry ice, and lymph nodes (LNs) were pre-fixed in 4% Paraformaldehyde (PFA) and then embedded in Tissue-Tek^®^ O.C.T. Compound (VWR) for cryo-sectioning as described below.

### Flow Cytometry

Spleens and LNs were harvested into ice-cold MACS buffer (1x PBS + 2% FCS + 1 mM HEPES) and mechanically dissociated using pestles in 1.5 mL Eppendorf tubes. Samples were then filtered through 70-µm cell strainers and centrifuged at 1500 rpm at 4 °C for 5 min. LN cell pellets were resuspended in MACS buffer. Spleen cell pellets were resuspended in RBC lysis buffer (155 mM NH_4_Cl, 12 mM NaHCO_3_, 0.1 mM EDTA), incubated for 5 min at RT and centrifuged as before. Splenocyte pellets were washed once with MACS buffer, centrifuged, and resuspended in MACS buffer. Single-cell suspensions were added to wells of 96-well round-bottom plates, centrifuged, and resuspended in 50 µL staining mix (appropriate Ab and viability dye at indicated concentrations, antibodies 1:200–1:400 and viability dye 1:1000, in MACS buffer). Fc receptors were blocked with anti-CD16/32 (clone 93) before staining. Samples were incubated for 30 min at 4 °C, followed by addition of 150 µL MACS buffer. Plates were subsequently centrifuged at 1500 rpm for 5 min, and supernatant was flicked out of the plates.

Flow cytometric analyses were performed on, an 8-color FACSCanto2 (BD Biosciences) and a 16-color LSRII (BD Biosciences), and Cytek Aurora spectral flow cytometer depending on the experiment. Data were analyzed with FlowJo software, v10.1).

### Immunofluorescence confocal microscopy and image analysis

Fresh spleen tissue was embedded in Tissue-Tek O.C.T. Compound (VWR), frozen on dry ice and stored at −80 °C until sectioning. Fresh frozen spleens were cut into 12-16 μm sections and air dried for 30 minutes, after fixing in 4% PFA for 10 min at RT, sections were permeabilized with 0.2% Triton X-100 in PBS for 10 min at RT and washed in PBS. After blocking for 1 hr at RT with blocking buffer (PBS, 0.01% Tween20, 2% BSA, and 3% Fetal Bovine Serum (FBS), sections were stained O.N. at 4 °C with antibodies (Table 2) diluted in blocking buffer. Slides were washed three times in PBS, 0.01% Tween20 and mounted with FluoroGel (Electron Microscopy Sciences) prior to image acquisition. Images were acquired with an OLYMPUS FV3000R resonant scanning confocal microscope equipped with four laser lines (405, 488, 514, and 633 nm), hybrid galvo and fast resonant scanning capabilities, ultra-sensitive GaAsP detectors with full spectral imaging and a motorized XYZ stage for tiling. For higher resolution, images were also acquired with 30X and 60X silicone oil objective and stitched using Olympus cell Sens software. The images were processed in Fiji.

### Autoantibody measurement

#### HEp-2 Immunofluorescence assay

Antinuclear antibodies in serum samples were analyzed by staining HEp-2 cells on 12-well HEp-2 slides (Fisher Scientific, cat. AN1012). In brief, 30 μL of diluted serum samples (1:20 dilution in 1X PBS) were applied to each well and incubated for 1 h at room temperature. After washing off excess serum, slides were washed twice with blocking buffer (0.1% Tween-20, 0.5% BSA in 1X PBS), 5 min each. Slides were then incubated in a secondary antibody mixture (anti-IgG2c-AF488 and Cholera Toxin B-AF647) in blocking buffer at room temperature for 1 hr to detect anti-nuclear antibodies. Following 2 washes with blocking buffer, DAPI Fluoromount-G mounting medium (Southern Biotech, cat# 0100-20) was added to slides and coverslips were applied. Images were acquired with an Olympus FV3000R resonant scanning confocal microscope with a 100x silicone oil objective.

#### Anti-single-stranded DNA autoantibody measurement by ELISA

ssDNA salmon DNA (20 μl/ml; D9156, Sigma-Aldrich) was diluted in 5% dimethyl sulfoxide in carbonate/bicarbonate coating buffer and incubated at 95°C for 10 min. It was then coated onto a 96-well ELISA plate and incubated overnight. The plate was washed three times with wash buffer (0.01% Tween 20 in PBS), then blocking buffer [5% bovine serum albumin (BSA) and 1% fetal calf serum in PBS] was added at 37°C for 1 hour. The plate was washed three times with wash buffer, and sera [diluted 1:20 in 10% blocking buffer in PBS (dilution buffer)] were added to the plate and incubated at 4°C overnight. Cell culture supernatants were added undiluted. The plate was washed three times, goat anti-mouse IgG2c-HRP (1079-05, Southern Biotech) was diluted in dilution buffer, and the plate was incubated at 37°C for 1 hour. After washing the plate three times, TMB substrate was added for 15 to 20 min, and the reaction was stopped with H_2_SO_4_. The 450-nm absorbance was measured on the xMark microplate spectrophotometer (Bio-Rad). C11 (564Igi Ab derived from a hybridoma) was used to standardize absolute levels of anti-ssDNA Ab.

### Statistical analysis

Two-tailed Mann–Whitney tests, one-way analyses of variance with Tukey correction and one-tailed unpaired Student’s t-tests were performed with Prism 9 (GraphPad).

## Acknowledgements

We thank Drs. V. Michael Holers (Univ Colorado School of Medicine, Denver) and Bridget Huber (Tufts New England School of Medicine) for providing the human CD21 BAC transgenic mice bred onto the murine mCD21^-/-^background; Dr. Theresa Imanishi-Kari (Tufts New England School of Medicine) for the 564 Igi strain, and Dr. T. Kurosaki (RIKEN-Yokohama) for the S1PR2-tdTom BAC transgenic mice.

## Author contributions

All authors contributed to the design of the research; MCC supervised the project. M.C.C and A.R conceived the project. M.C.C and A.R designed and planned all the experiments. AR performed most of the experiments with assistance from D.Z and K.O. A.R did data analysis/interpretation and wrote the paper. All authors provided feedback on manuscript.

## Funding

This study was supported by the National Institutes of Health (NIH grant 5R01AR074105), the Lupus Research Alliance (LRA grant #1149175), and the Department of Defense (DoD grant #HT9425-24-1-0598).

## Competing interests

M.C.C. is a co-inventor on U.S. Patent Application US20240376225A1, “CD21 antibodies and uses thereof,” filed by The Children’s Medical Center Corporation and Pfizer Inc., which covers the anti-CD21 antibody (clone 658) used in this study. The antibody was provided through a research collaboration with Pfizer Inc. The remaining authors declare no competing interests.

---

**Fig s1.**
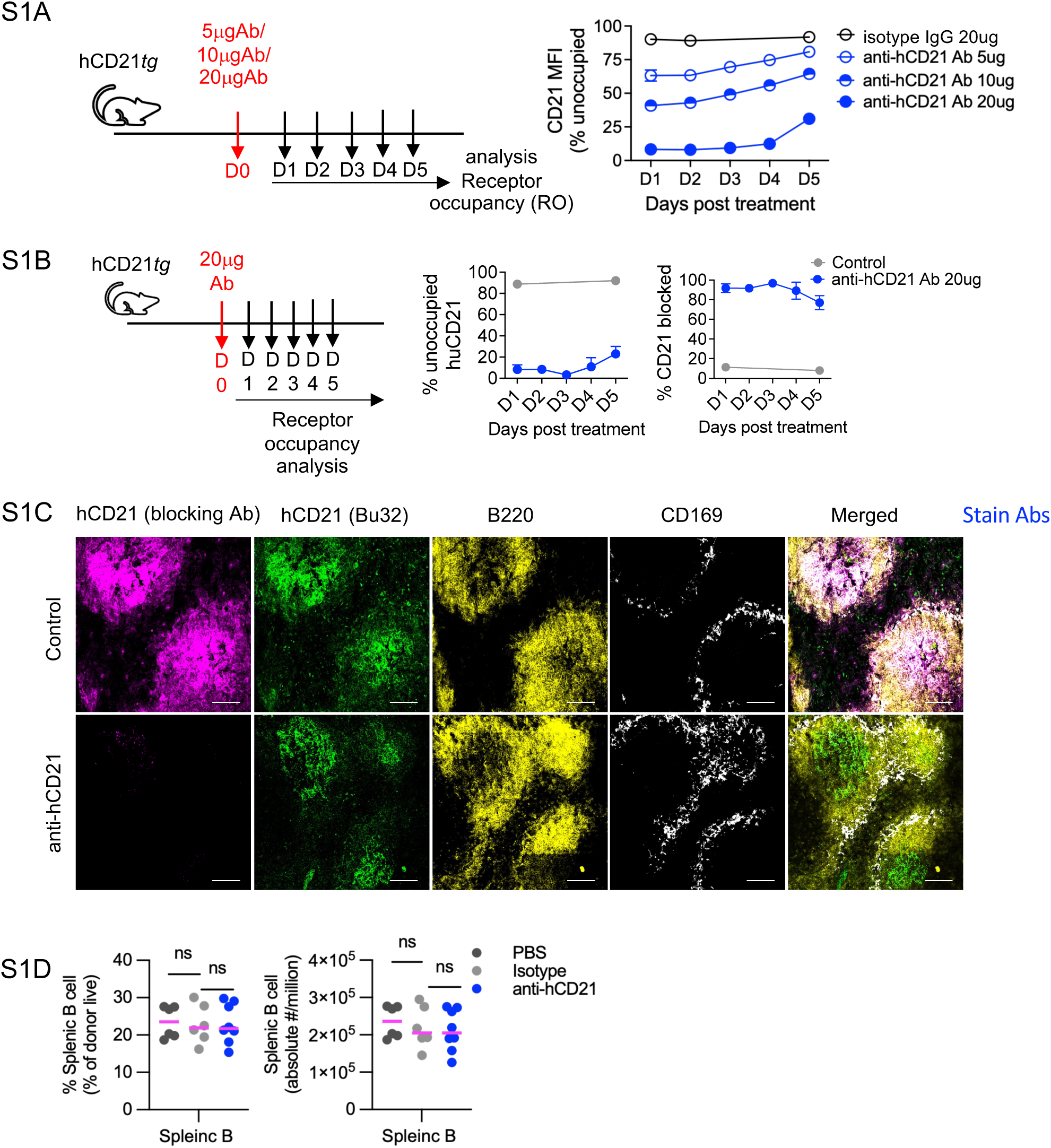
A) Dose-titration experimental design: hCD21*tg* mice received a single intraperitoneal injection of isotype control IgG (20 μg) or anti-hCD21 antibody (5, 10, or 20 μg), with CD21 receptor occupancy (% unoccupied, by MFI) assessed daily for 5 days. **B)** Repeated-timepoint receptor occupancy assay following a single 20 μg anti-hCD21 dose: percent unoccupied huCD21 and percent CD21 blocked, control versus anti-hCD21 antibody, assessed daily for 5 days. **C)** *In vivo* blocking efficiency of anti-hCD21 antibody (FDC blockade). Spleen sections from B6 WT + 564Igi → irradiated hCD21*tg* bone marrow chimeras. Donor-derived B cells do not express hCD21; FDCs are radioresistant and host-derived, so follicular hCD21 signal is FDC-specific. Mice received isotype control or anti-hCD21. Spleen sections were stained *ex vivo* with conjugated blocking anti-hCD21 (magenta; loss of signal indicates epitope occupancy by antibody given *in vivo*), non-competing clone Bu32 (green; total surface hCD21), B220 (yellow), and CD169 (white). Blocking efficiency of anti-hCD21 antibody ranged from 85–97%. Scale bars = 100 μm. **D)** absolute cell counts and frequence of splenic B cell numbers in anti-hCD21 antibody-, isotype control-, and PBS-treated mice. Scale bars, 100 µm. Data are mean ± SEM, pooled from 2 independent experiments (n = 6 PBS, 6 control, 8 anti-hCD21-treated mice). Each symbol represents one mouse. Statistical significance was determined by unpaired two-tailed Welch’s t test (or Mann–Whitney U test). ns, not significant.

**Fig. S2.**
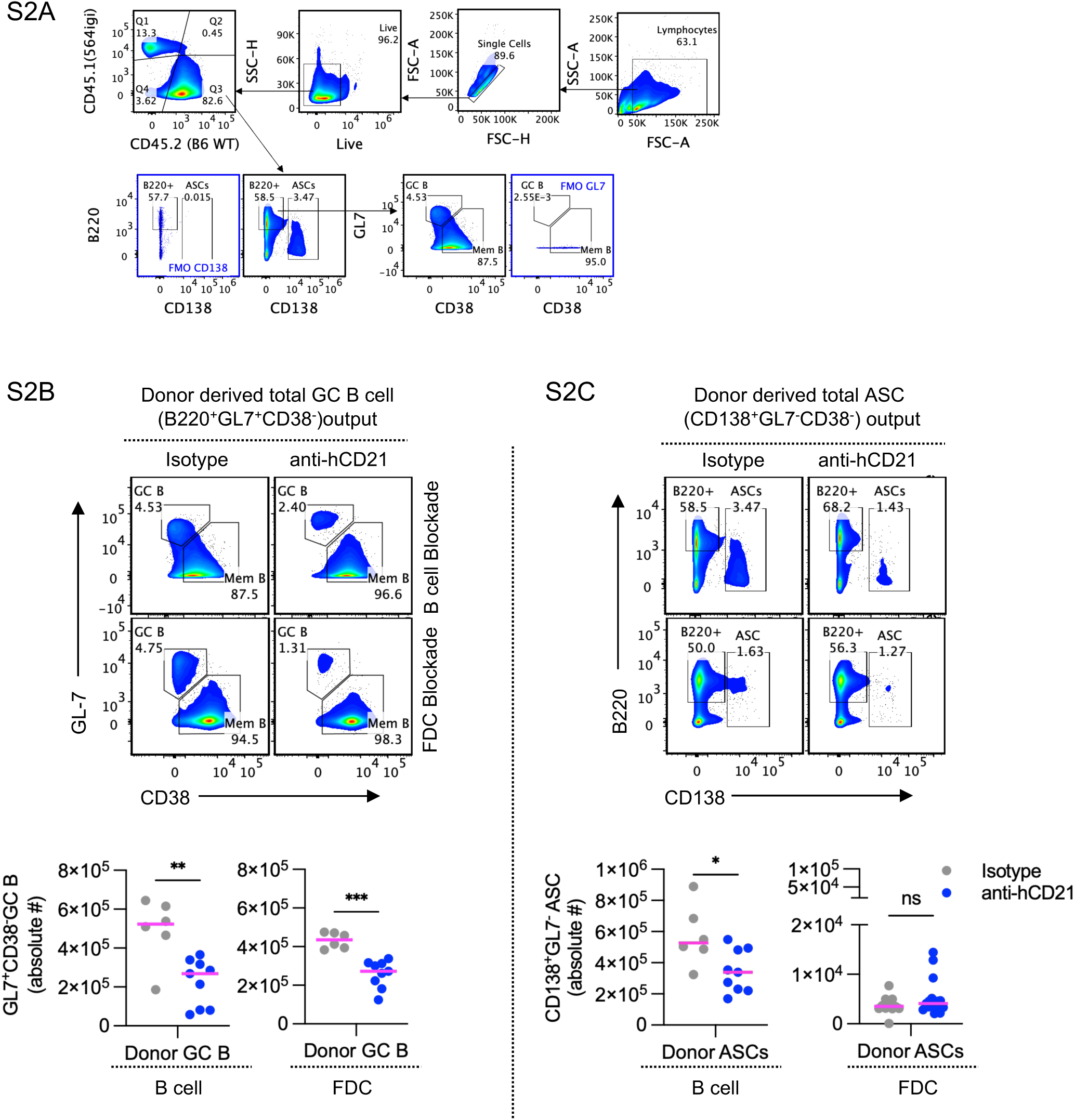
Gating strategy and quantification of donor-derived GC B cell and ASC responses in B cell- and FDC-blockade chimeras. **A)** Hierarchical flow cytometry gating strategy used to identify donor-derived (CD45.2) total GC B cells (B220^+^GL7^+^CD38^-^) and ASCs (B220^+^CD138^+^). **B)** Representative flow plots and absolute cell counts of donor-derived total GC B cells (GL7^+^CD38^-^) and **C)** ASCs (CD138^+^GL7^-^) in B cell-blockade and FDC-blockade chimeras, isotype control versus anti-hCD21 Ab. Data are mean ± SEM, pooled from 2 independent experiments (n = 6 control, 9 anti-hCD21-treated mice). Each symbol represents one mouse. Statistical significance was determined by unpaired two-tailed Welch’s t test (or Mann–Whitney U test); *P < 0.05, **P < 0.01, ***P < 0.001. *ns*, not significant.

**Fig. S3.**
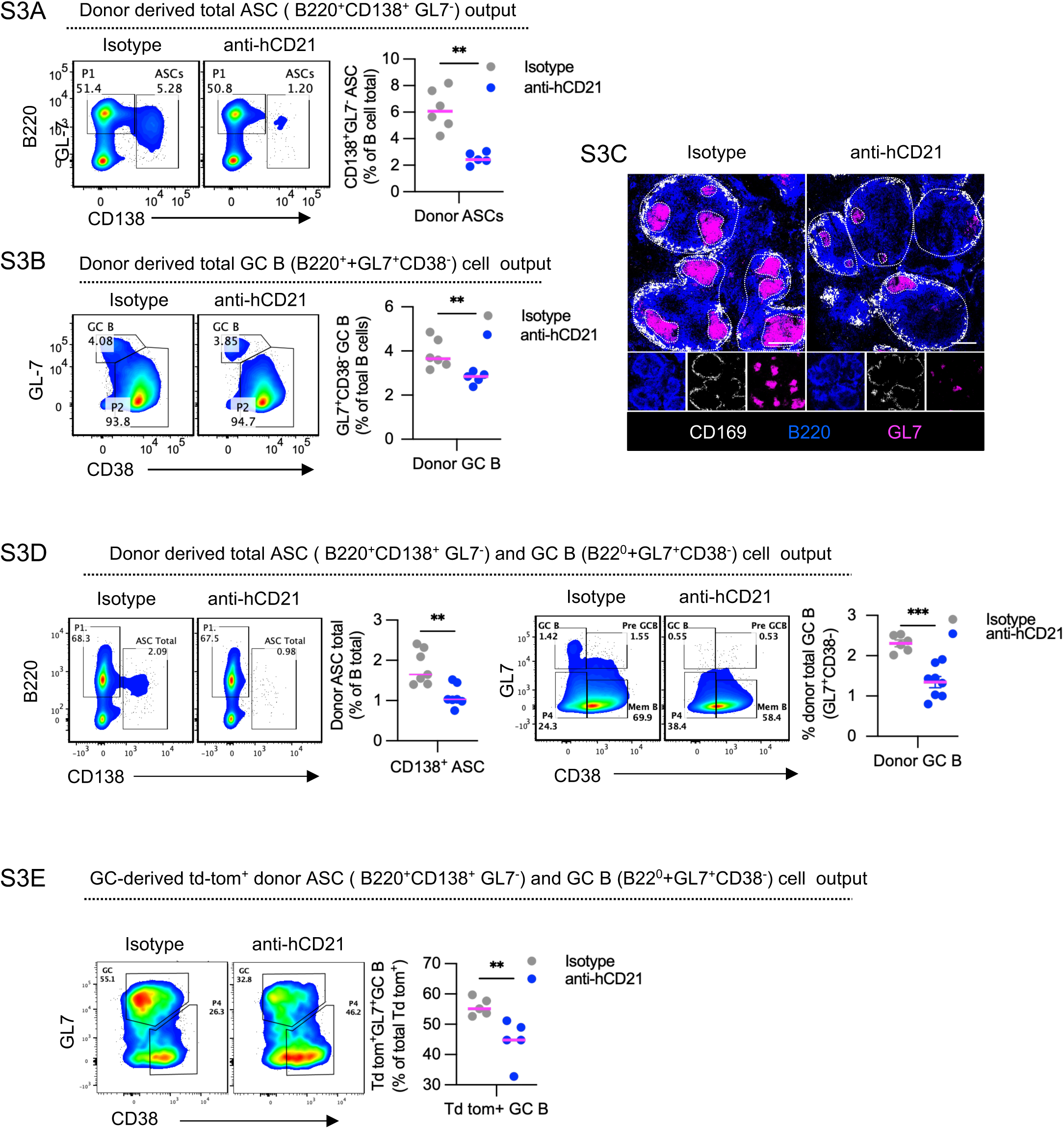
Dual CD21 blockade synergistically reduces donor-derived GC B cell and ASC output, including in the extended-treatment cohort. A-C) Dual blockade, 3 antibody injections. A) Representative flow plots and quantification of donor-derived total ASCs (B220^+^CD138^+^GL7^-^), isotype control versus anti-hCD21 Ab. B) Representative flow plots and quantification of donor-derived total GC B cells (B220^+^GL7^+^CD38^-^). C) Representative immunofluorescence (CD169, B220, GL7) of splenic follicles, control versus anti-hCD21 Ab. Scale bars = 100 μm. D-E) 5-injection dual blockade; D) Representative flow plots and quantification of donor-derived total ASC: *left panel*, and GC B cell output; *right panel*, following extended antibody treatment. E) Representative flow plots and quantification of GC-derived (Td-tomato^+^) donor GC B cell output following extended dual blockade. Data are mean ± SEM, pooled from 2 independent experiments. Each symbol represents one mouse. Statistical significance was determined by unpaired two-tailed Welch’s t test (or Mann–Whitney U test, Prism v10.1); **P < 0.01, ***P < 0.001. *ns*, not significant.

**Fig. S4.**
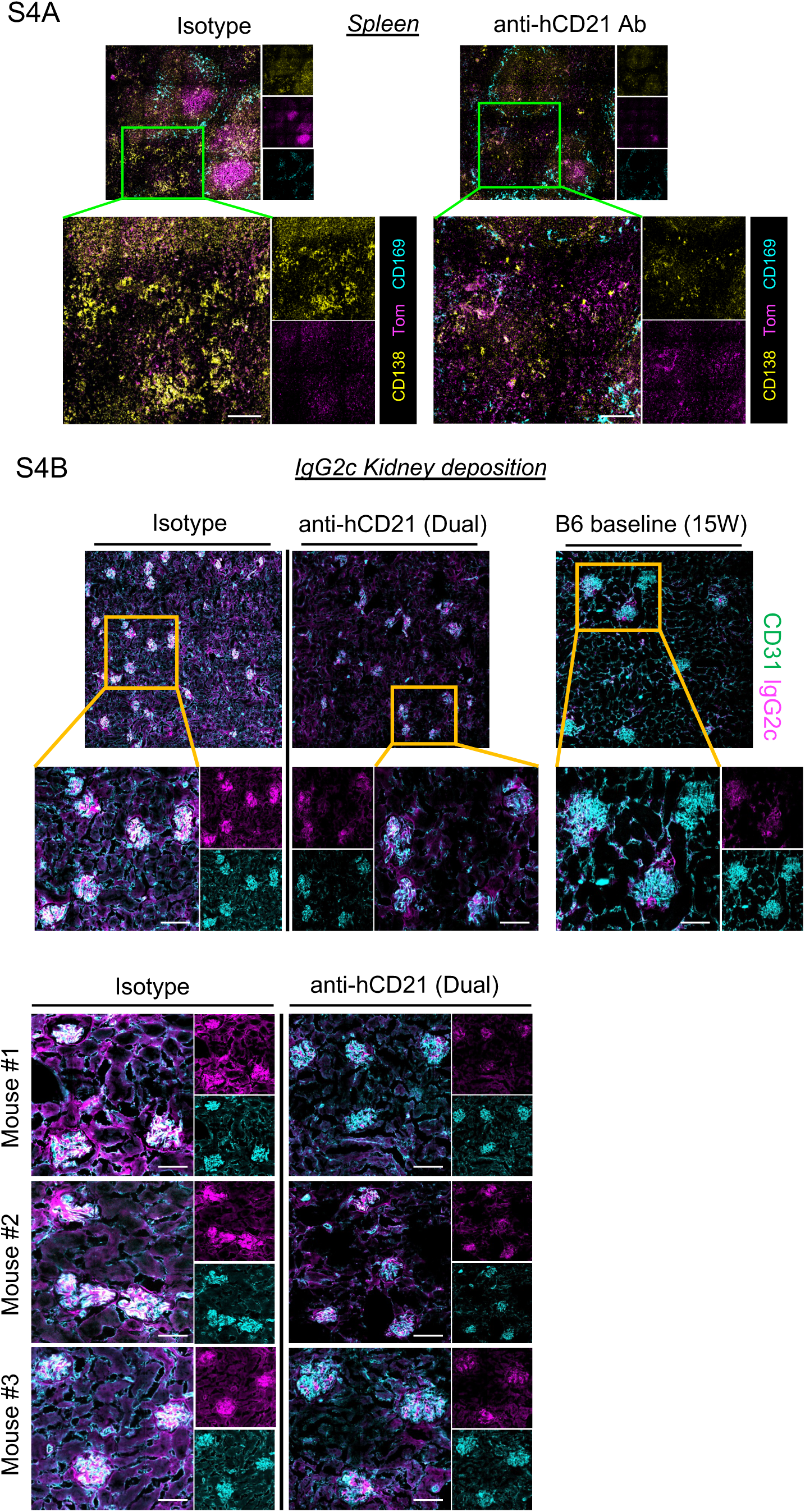
Splenic and renal immune-complex deposition following extended dual CD21 blockade. **A)** Representative immunofluorescence of spleen sections (CD138, tdTomato, CD169), in isotype control versus anti-hCD21 Ab, showing total CD138+ ASC signal (yellow) in red pulp. **B)** Representative immunofluorescence of kidney sections (CD31, IgG2c) from isotype control vs. anti-hCD21 Ab (dual)-treated, and 15-week-old B6 baseline mice; ***upper panel***, additional individual-mouse replicates, illustrating the synergistic decrease in glomerular IgG2c deposition with prolonged dual blockade in in isotype control versus anti-hCD21 Ab treaded animals . Scale bars = 100 μm.

**Fig. S5.**
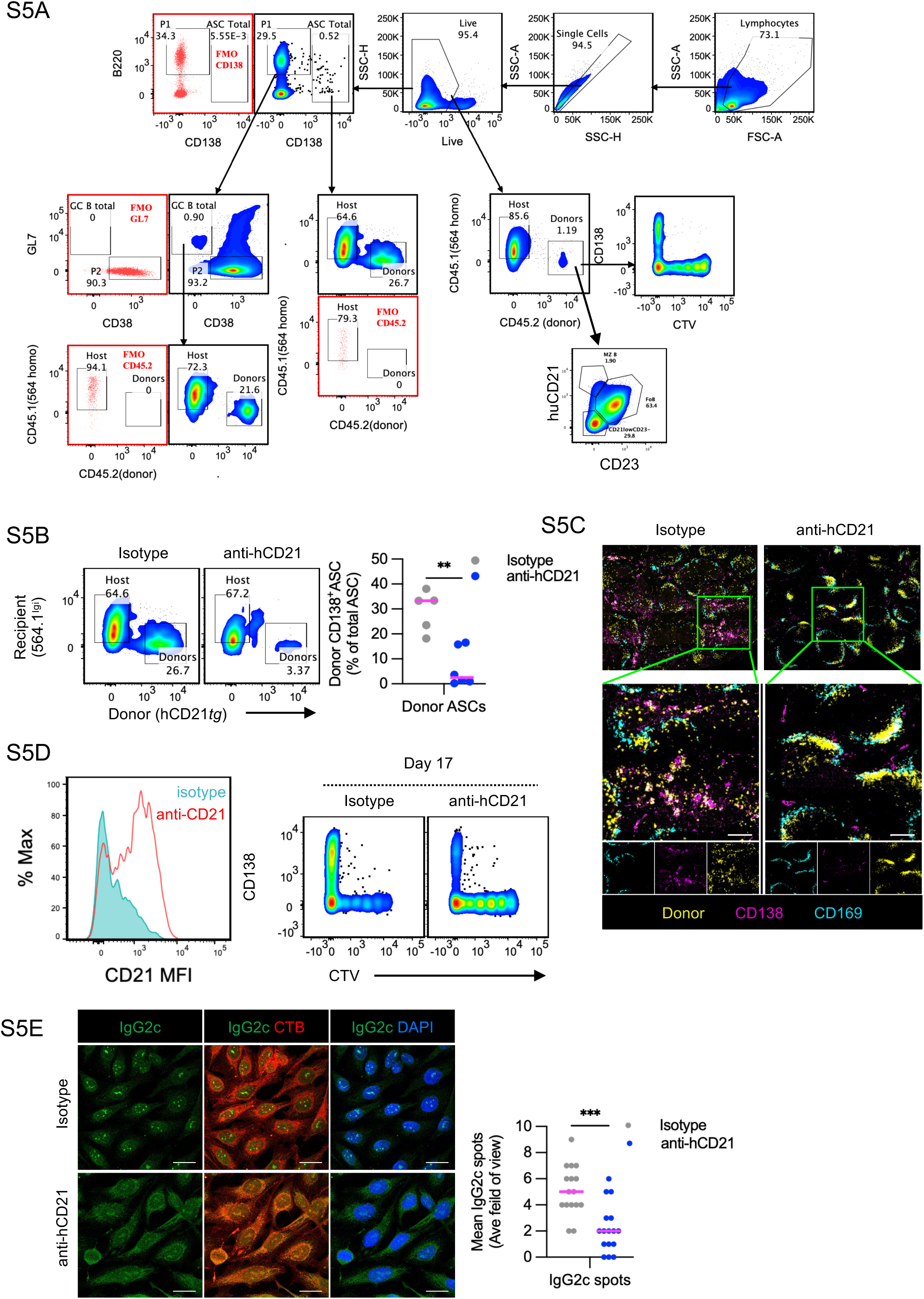
CD21 blockade halts activation of naïve and recently activated B cells transferred into an ongoing autoreactive response. A) Gating strategy for donor (CD45.2, hCD21*tg*) versus recipient (CD45.1, 564.1Igi) B cells, GC B cells, ASCs, and follicular/activated (hCD21 versus CD23) subsets. **B) B)** Representative flow plots and quantification of donor-derived CD138^+^ ASCs (recipient 564.1Igi versus donor hCD21*tg*), isotype control versus anti-hCD21 Ab at D17. C) Representative immunofluorescence (donor, CD138, CD169) showing donor tdTom^+^ B cell/ASC localization, in the splenic red pulp, isotype control versus anti-hCD21 Ab. Scale bars = 200 μm. D) Representative CD21 expression histogram (isotype control versus anti-hCD21 Ab) and flow plots (CD138 versus CTV) at D17 showing less generation CD138^+^ cells and reduced CTV dilution. E) 1:30 dilution of serum from isotype control versus anti-hCD21 Ab animals was applied onto HEp-2 slides to visualize donor derived (IgG2c) antibody reactivity to auto-antigens. Representative HEp-2 cell immunofluorescence (IgG2c, IgG2c+CTB, IgG2c+DAPI) and quantification of IgG2c spot counts, isotype control versus anti-hCD21 Ab. Images taken at 100x magnification, scale bars = 20 μm. Data represent mean ± SEM for 2 independent experiments. ***p < 0.001 (Mann–Whitney U, paired t test, Prism v10.1).

**Fig. S6.**
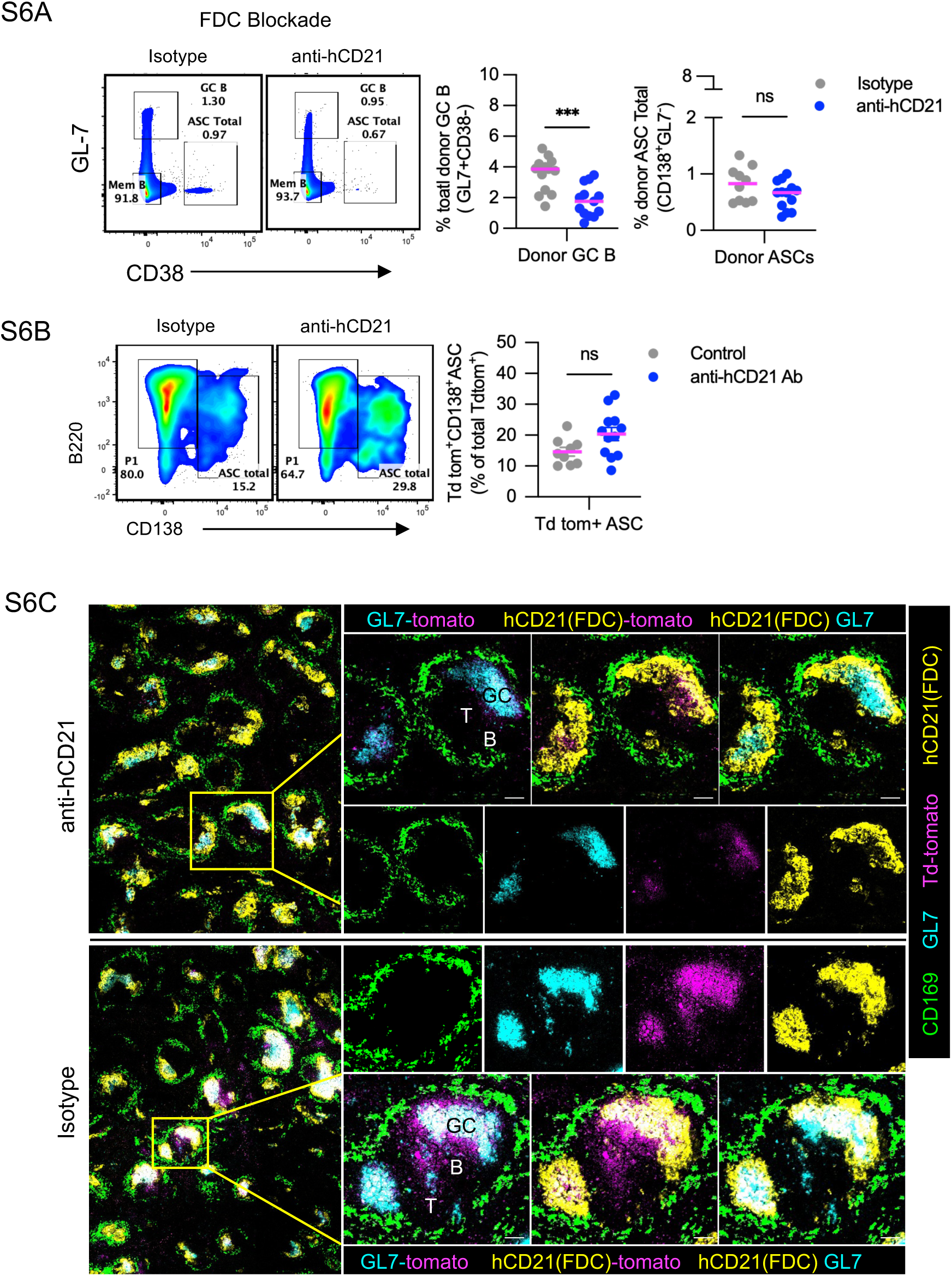
FDC-specific CD21 blockade disrupts GC B cell–FDC engagement. **A)** Representative flow plots and quantification of total donor GC B cells and ASCs following FDC-specific blockade, isotype control versus anti-hCD21 Ab. **B)** Representative flow plots and quantification of GC-derived tdTom^+^ donor ASCs (GL7^-^CD138^+^) following FDC-specific blockade, isotype control versus anti-hCD21 Ab. **C)** Representative immunofluorescence (GL7-tdTomato, hCD21[FDC]–tdTomato, hCD21[FDC]-GL7) showing loss of GC B cell–FDC colocalization within the T/B/GC zones, isotype control versus anti-hCD21 Ab. Scale bars = 100 μm. Data represent mean ± SEM for pool of 2 independent experiments. ***p < 0.001 (Mann–Whitney U, paired t test, Prism v10.1).

**Fig. S7.**
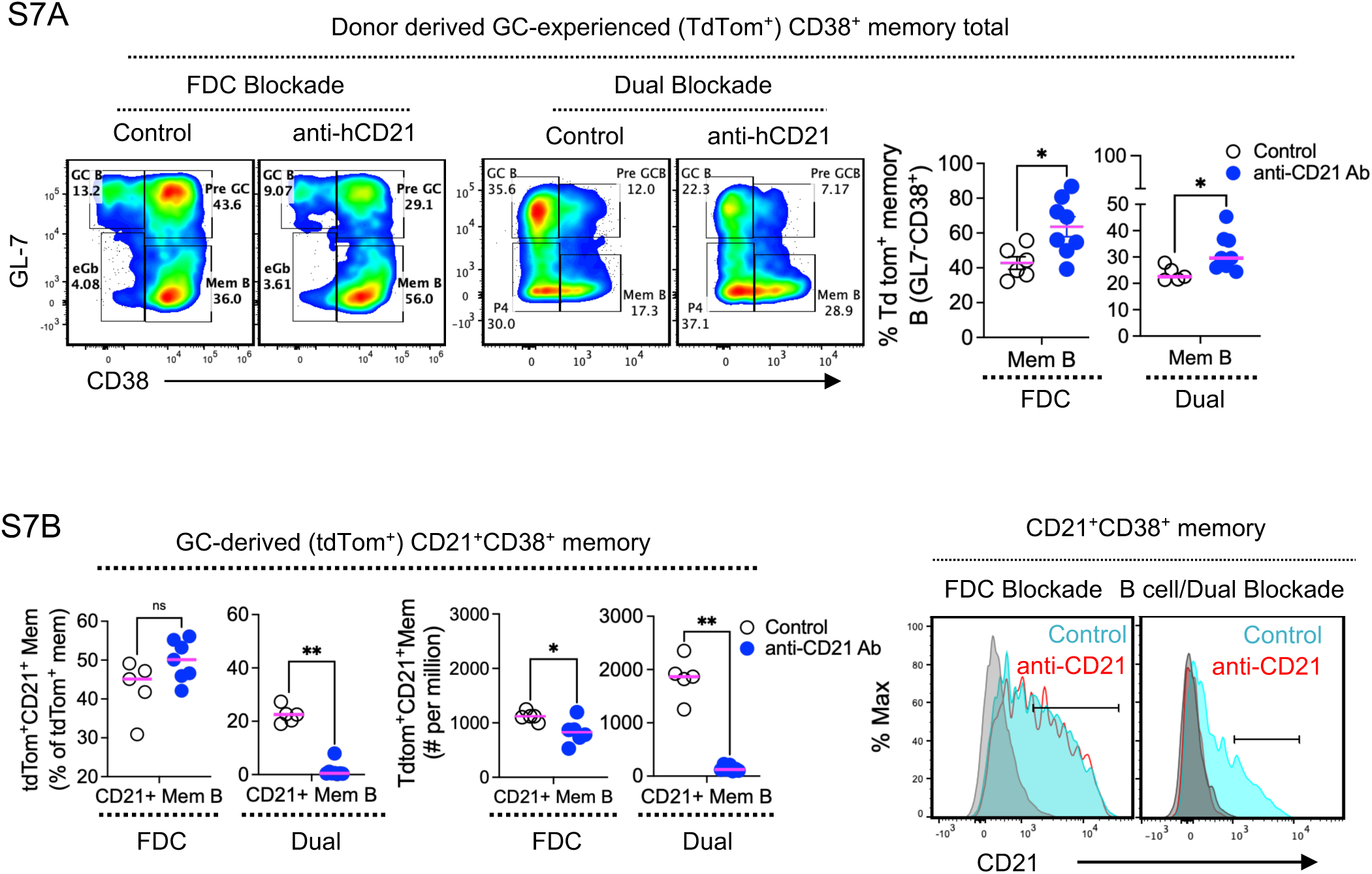
Differential effects of FDC-versus dual-blockade on the GC-experienced memory B cell compartment. A) Representative flow plots and quantification of GC-experienced (tdTomato^+^) CD38^+^ memory B cells following FDC blockade and dual blockade, isotype control versus anti-hCD21 Ab. B) Quantification of GC-derived (tdTomato^+^) CD21^+^CD38^+^ memory B cell frequency and absolute number; ***left panel***, and representative CD21 expression histograms; ***right panel***, for FDC-blockade and dual-blockade cohorts. Data represent mean ± SEM for pool of 2 independent experiments. *p < 0.05, **p < 0.01 (Mann–Whitney U, Prism v10.1). *ns*, not significant.

## Notes

### Competing Interest Statement

The authors have declared no competing interest.

