## Supplemental tables for "Dual cellular targeting of complement receptor 2 (CR2/CD21) synergistically suppresses autoreactive germinal center activity and autoimmunity in lupus"

Table 1: Genotyping primers

| **S.No.** | **Gene** | **Primer Sequence** |
| --- | --- | --- |
| 1 | *hCD21* | Fwd: 5'-GTT TTC TTG GCT CTC GTC GC-3' |
|  |  | Rev: 5'-TCG GGT GTA CAG AAA CAG CC-3' |
| 2 | *mCD21* | WT Fwd: 5'-TGT CAG GCT CCT CCT AAA ATT ATC-3' |
|  |  | WT rev: 5'-CTT TAC AAA GAC GGA TTT CTA TA-3' |
|  |  | Mut rev 5'-CCA CTG TCC TTT CCT AAT AAA ATG-3' |
| 3 | *S1PR2^CreER^* | Fwd: 5'-GCG GTC TGG CAG TAA AAA CTA TC-3' |
|  |  | Rev: 5'-CAG CTA CAC CAG AGA GAC GGA AAT C-3' |
| 4 | *TdTomato* | WT Ford: 5'-AAG GGA GCT GCA GTG GAG TA-3' |
|  |  | WT Rev: 5'-CCG AAA ATC TGT GGG AAG TC-3' |
|  |  | Mutant Rev: 5'-GGC ATT AAA GCA GCG TAT CC-3' |
|  |  | Mutant Fwd: 5'-CTG TTC CTG TAC GGC ATG G-3' |
| 5 | *CxCL13^Cre^* | Cre Fwd: 5'-GCTGCCTGGTGCAAGCTGAA-3' |
|  |  | Cre Rev: 5'-ACCATCGGTGCGGGAGATGT-3' |
| 6 | *iDTR* | Common Fwd: AAA GTC GCT CTG AGT TGT TAT |
|  |  | Common Rev: GGA GCG GGA GAA ATG GAT ATG |
|  |  | Mut Rev: GCG AAG AGT TTG TCC TCA ACC |

Table 2: Antibodies and reagents.

| ***S. No*** | ***Target*** | ***Clone/***  ***Catalogue^#^*** | ***Company*** | ***Fluorochrome (Flow, IF)*** |
| --- | --- | --- | --- | --- |
| 1 | B220 | RA3-6B2 | Biolegend | Buv395, BV711, pacific blue, APC, |
| 2 | CD19 | 6D5 | Biolegend | AF700, PE/Cy7 |
| 3 | CD45.1 | A20 | Biolegend | BV605, pacific blue, APC |
| 4 | CD45.2 | 104 | Biolegend | PerCP/Cy5.5, BV650, PE/Cy7, AF488 |
| 5 | IgD | 11-26c.2a | Biolegend | Pacific blue, PE/Cy7, FITC |
| 6 | IgM | RMM-1 | Biolegend | BV605, PE, PerCP/Cy5.5 |
| 7 | hCD21 | Bu32 | Biolegend | PE, FITC |
| 8 | hCD21 | 658 | Pfizer | AF647 |
| 9 | mCD21 (CD21/35) | 7E9 | Biolegend | BV421, BV711 |
| 10 | GL7 | GL7 | Biolegend | AF488, APC, pacific blue, PerCP/Cy5.5 |
| 11 | CD138 | 281-2 | Biolegend | BV510, APC |
| 12 | CD23 | B3B4 | Biolegend | PE/Cy7, BV786, BV510 |
| 13 | FCRL5 | 6767G^#^ | R&D systems | FITC, APC |
| 14 | CD38 | 90 | Biolegend | PE/Cy7, AF700 |
| 15 | CD11c | N418 | Biolegend | PE/Cy7, PerCP/Cy5.5 |
| 16 | Human IgG1/κ Isotype | 569605^#^ | BD Bioscience | Unconjugated |
| 17 | mCD21 | 7G6 | BD Bioscience | Blocking unconjugated |
| 18 | CD3e | 17A2 | Biolegend | APC, FITC |
| 19 | CD4 | RM4-5 | Biolegend | APC, FITC, pacific blue |
| 20 | CD45 | 30-F11 | Biolegend | AF488, APC, pacific blue, PerCP |
| 21 | CD80 | 16-10A1 | Biolegend | Alexa flour 700, APC |
| 22 | CD16/32 | 93 | Biolegend | Unconjugated |
| 23 | CD11b | M1/70 | Biolegend | AF700, PE/Cy7, PE |
| 24 | CD169 | 3D6.112 | Biolegend | BV421, FITC, APC |
| 25 | eflour780 | 65-0865-14^#^ | eBioscience | Fixable viability dye |
| 26 | Goat anti-mouse IgG2c | 1079-30^#^ | Southern Biotech | AF488 |
| 27 | HEp-2 slides (12-well) | AN1012^#^ | Fisher Scientific | - |
| 28 | Fixation buffer | 420801^#^ | Biolegend | - |
| 29 | MojoSort Mouse Pan B Cell Isolation Kit | 480088^#^ | Biolegend | - |
| 30 | MACS LS columns | 130-042-401^#^ | Miltenyi Biotec | - |
| 31 | DAPI Fluoromount-G | 0100-20^#^ | Southern Biotech | - |
| 32 | Tissue-Tek O.C.T. compound | 25608-930^#^ | VWR | - |
| 33 | 16% Paraformaldehyde | 15710-S^#^ | Electron Microscopy Sciences | - |
| 34 | Super Frost Plus slides | 12-550-15^#^ | Fisher Scientific | - |
| 35 | 70 µm cell strainers | 22-363-548^#^ | Fisher Scientific | - |
| 36 | HBSS | 21-022-CV^#^ | Corning | - |
| 37 | RPMI 1640 with L-glutamine | 10-040-CV^#^ | Corning | - |
| 38 | HEPES, 1 M | 25-060-CI^#^ | Corning | - |
| 39 | BSA, heat-shock treated | BP1600-100^#^ | Fisher Scientific | - |
| 40 | Fetal bovine serum | F0926-500ML^#^ | Sigma-Aldrich | - |
|  | β-Mercaptoethanol | 21985-023^#^ | Thermo Fisher | - |

^#^Catalouge number.
